# The structural basis of odorant recognition in insect olfactory receptors

**DOI:** 10.1101/2021.01.24.427933

**Authors:** Josefina del Mármol, Mackenzie Yedlin, Vanessa Ruta

## Abstract

Olfactory systems must detect and discriminate an enormous diversity of chemicals in the environment. To contend with this challenge, diverse species have converged on a common strategy in which odorant identity is encoded through the combinatorial activation of large families of olfactory receptors (ORs), thus allowing a finite number of receptors to detect an almost infinite chemical world. Although most individual ORs are sensitive to a variety of odorants, the structural basis for such flexible chemical recognition remains unknown. Here, we combine cryo-electron microscopy with functional studies of receptor tuning to gain insight into the structural and mechanistic basis of promiscuous odorant recognition. We show that OR5 from the jumping bristletail, *Machilis hrabei*, assembles as a homo-tetrameric odorant-gated ion channel with broad chemical tuning. We elucidated the structure of OR5 in multiple gating states, alone and in complex with two of its agonists—the odorant eugenol and the insect repellent DEET. Both ligands bind to a common binding site located in the transmembrane region of each subunit, composed of a simple geometric arrangement of aromatic and hydrophobic residues. We reveal that binding is mediated by hydrophobic, non-directional interactions with residues distributed throughout the binding pocket, enabling the flexible recognition of structurally distinct odorants. Mutation of individual residues lining the binding pocket predictably altered OR5’s sensitivity to eugenol and DEET and broadly reconfigured the receptor’s tuning, supporting a model in which diverse odorants share the same structural determinants for binding. Together, these studies provide structural insight into odorant detection, shedding light onto the molecular recognition mechanisms that ultimately endow the olfactory system with its immense discriminatory capacity.

## Introduction

The olfactory system faces a unique challenge amongst sensory modalities due to the inordinate complexity of the chemical world. While light waves vary continuously along a single axis in amplitude and frequency, odorants differ discretely along an enormous number of dimensions in their molecular structure and physicochemical properties. Whereas just three photoreceptors are sufficient to sense the entire spectrum of visible light, large repertoires of olfactory receptors appear necessary to detect and discriminate amongst the diversity of chemicals in the environment^1–3^. In mammals and other vertebrates, odor detection is mediated by G-protein coupled receptors that signal through canonical second-messenger cascades^4,5^. In contrast, insects detect volatile chemicals via a unique class of odorant-gated ion channels^6–8^ comprised of two subunits: a highly conserved coreceptor (Orco) subunit^9,10^ and a highly divergent odorant receptor (OR) subunit that harbors the odorant-binding site and confers chemical sensitivity to the heteromeric complex. Olfactory receptors have massively expanded and diversified across insect lineages to emerge as possibly the largest and most divergent family of ion channels in nature^2^, with potentially millions of distinct variants distributed across hundreds of thousands of different insect species. The rapid evolution of ORs is thought to facilitate the chemosensory specialization of insects, endowing each species with the ability to detect chemicals relevant to its ecological niche^11^.

While mammals and insects rely on distinct molecular mechanisms for olfactory detection, they share a common neural logic for olfactory perception in which the identity of an individual odorant is encoded through the combinatorial activation of a unique ensemble of olfactory receptors and associated sensory neurons^1,3,12^. Central to this sensory strategy is that most ORs recognize a variety of structurally diverse ligands and display overlapping molecular receptive fields^13–16^. Combinatorial coding is thought to endow the olfactory system with its immense discriminatory power, expanding its potential coding capacity from the number of receptors within the genome to the number of combinations among them. Yet how a single OR can promiscuously detect different odorants has remained elusive, in part due to the lack of explicit structural models for odorant recognition. One possibility is that ORs possess multiple odorant-binding sites that differ in their chemical specificity^17^, thus expanding the receptor’s tuning breadth. Alternatively, a single binding site may recognize just one chemical feature of an odorant, rendering the same receptor sensitive to a variety of ligands that share this common moiety^18,19^. Completely orthogonal mechanisms have also been proposed to explain receptor tuning, including the hypothesis that ORs recognize the vibrational modes of odorants rather than their stereochemical features^20,21^. Thus, while the physicochemical properties of an odorant are thought to be a key determinant of its olfactory percept^14,22–24^, almost nothing is known about how this is defined at the structural level.

Here, we leveraged the evolutionary diversification of insect ORs to shed light on the structural basis of odorant recognition. Neopteran insects, which encompass the staggeringly diverse clade of winged species, each express a distinct repertoire of ORs that vary both in number and sequence, along with a single almost invariant Orco^2,25^ (Fig. **1a**). Given Orco’s striking sequence conservation across insect lineages and its essential role in olfactory transduction^9,26–28^, it was previously proposed to represent the evolutionarily most ancient form of the insect olfactory receptor complex^25^. Indeed, in contrast to neopteran ORs, Orco can autonomously assemble into homo-tetrameric cation channels that, while insensitive to volatile odorants, can be activated by synthetic agonists^29^. We previously elucidated the structure of an Orco homo-tetramer^30^, offering initial insight into the architecture of the insect olfactory receptor family and how it accommodates the striking sequence diversity of ORs. However, given that Orco is nearly chemically inert, how odorant recognition is achieved or transduced to channel gating remains unclear.

**Figure 1.**
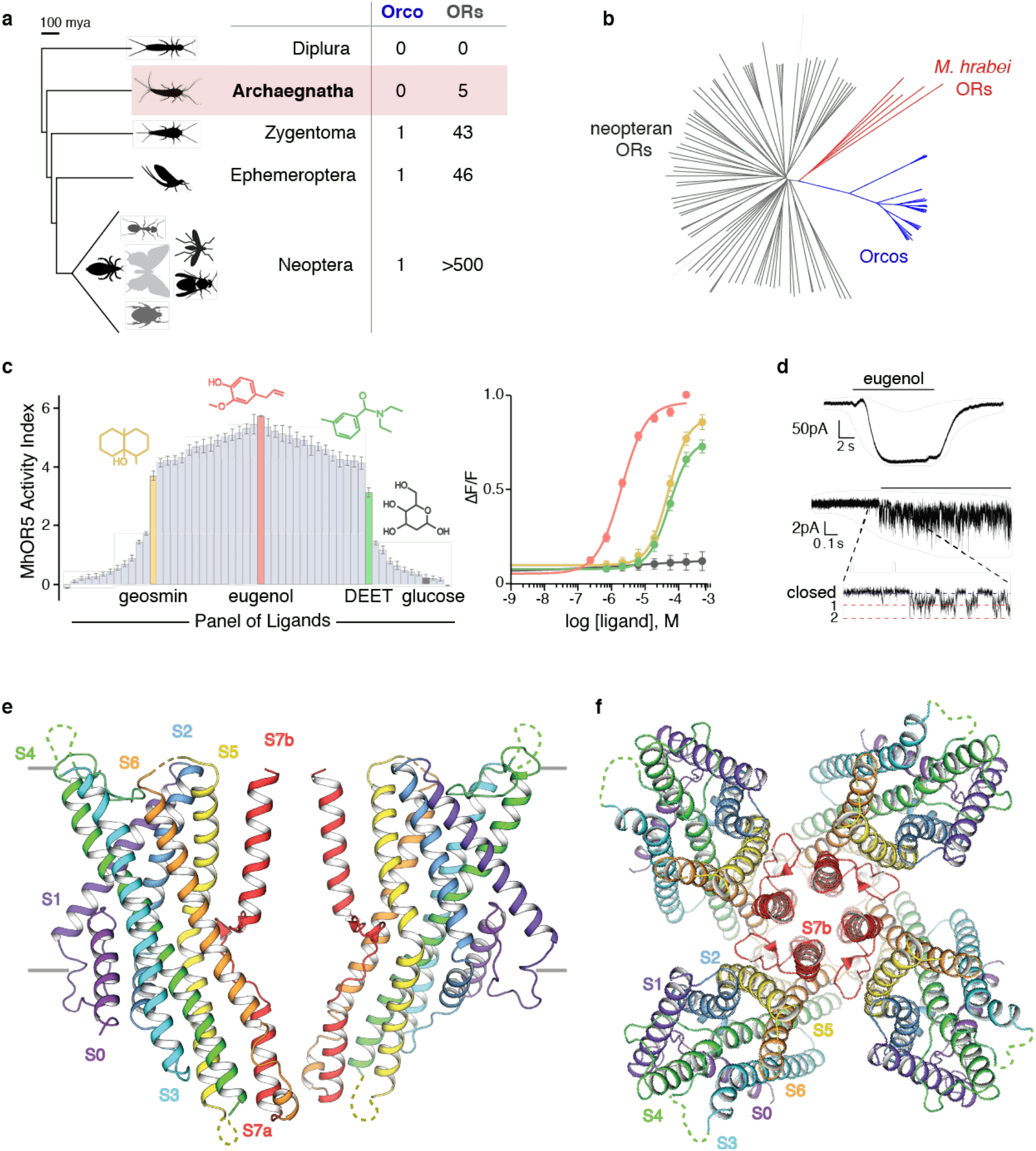
MhOR5 forms a homo-tetrameric ligand-gated ion channel activated by a broad panel of odorants. **a**. A phylogenetic tree of select insect clades and the number of OR and Orco genes present in their genomes. **b**. An unrooted tree of the insect OR gene family displaying the relationships between Orcos (blue), neopteran ORs (grey) and ORs from *M. hrabei* (red). Alignment was made from 28 Orco sequences, 82 OR sequences from 4 species (*Anopheles gambiae, Drosophila melanogaster, Nasonia vitripennis* and *Pediculus humanus*) using PROMALS3D (see Methods). **c**. Tuning curve of MhOR5 activity evoked by a panel of 54 ligands (left) (3 < n < 122; median n = 11.5). Dose-response curves of MhOR5 in the presence of eugenol (pink; n = 122), geosmin (gold; n = 11), DEET (green; n = 12), and glucose (grey; n = 4) (right). Additional data in Extended Data Fig. 2 and Extended Data Table 2. **d**. Electrophysiological recording from HEK cells expressing MhOR5. Eugenol-elicited inward currents in whole-cell recordings clamped at −80 mV (top) and single-channel recordings in outside-out patches clamped at −80 mV. **e, f**. Cryo-EM structure of MhOR5 shown from the side (**e**) and top (**f**). Each subunit is colored in rainbow palette from the N-term (purple) to the C-term (red). Two subunits are shown in the side view along with markers for the membrane in grey, while all four subunits are shown in the top view. Detailed description of metrics used in Methods.

Recent genomic analyses have revealed that some ancestral insects, such as the jumping bristletail, *Machilis hrabei*, possess a small number of putative OR genes but lack an apparent Orco ortholog^31^ (Fig. **1a**). This surprising observation raises the possibility that these primitive ORs may function as homomeric olfactory receptors, offering an entry point to explore the structural basis of odorant recognition in a biochemically simplified system. Here, we demonstrate that *M. hrabei* OR5 (MhOR5) forms a homo-tetrameric olfactory receptor with broad chemical tuning. Elucidating the structure of MhOR5, alone and in complex with two of its agonists, reveals how diverse odorants are recognized within a geometrically simple binding pocket through distributed hydrophobic interactions, suggesting a structural logic for this receptor’s promiscuous chemical tuning.

### *M. hrabei* OR5 is a broadly tuned odorant-gated ion channel

The *M. hrabei* genome harbors only five OR genes^31^. MhORs have been proposed to be amongst the most evolutionarily ancient members of the insect olfactory receptor family, arising prior to the emergence of Orco^31,32^. Despite sharing only 15-20% amino-acid conservation with either neopteran Orcos or ORs, MhORs display characteristics of both of these receptors in their pattern of amino-acid sequence conservation^32^. While little is known about the role these receptors play in chemosensory detection in the jumping bristletail, we reasoned that MhORs may form homomers, like Orco, that are gated by odorants, like ORs. To explore this possibility, we heterologously expressed each MhOR in HEK293 cells and found that MhOR1 and MhOR5 migrated with a similar pattern as Orco on non-denaturing native gels, indicating they assemble as tetramers (Extended Data Fig. **1a-c**). To assess whether these homomeric complexes function as chemoreceptors, we employed a high-throughput fluorescence assay^30^ in which we transiently transfected HEK cells with MhORs and the genetically-encoded calcium indicator GCaMP6s, enabling us to measure calcium flux in response to a panel of 54 small molecules over a range of concentrations. Our panel included an array of volatile odorants representing different chemical classes, including compounds known to be ecologically relevant for some insect species^15,16^, as well as several tastants and synthetic modulators of insect olfactory receptors. We found that MhOR5 was activated by many odorants but generally unresponsive to salts, sugars, or other tastants, consistent with a role in olfactory detection (Fig. **1c**, Extended Data Fig. **2**). MhOR5 was also activated by the insect repellent DEET and inhibited by the synthetic Orco agonist VUAA1^29^. As is characteristic of both insect and mammalian ORs^14–16^, many odorants activated MhOR5 with relatively low-affinity or sub-maximal efficacy, giving rise to heterogenous dose-response curves. Therefore, to quantitatively capture the complexity of odorant-evoked responses^33^, we defined an Activity Index for each odorant (-log(EC_50_) ∗ max ΔF/F) that reflects both its relative affinity and maximal efficacy (Extended Data Fig. **2a-c**). MhOR5 possessed a broad molecular receptive field analogous to many ‘generalist’ neopteran ORs^15,16^ (Extended Data Fig. **1e**) and responded to more than 68% of assayed odorants. In contrast, MhOR1 exhibited distinct and far more selective tuning, with strong responses evoked by only eight odorants from the same chemical panel (Extended Data Fig. **1d-f**). Both MhOR1 and MhOR5 were activated by ligands that spanned multiple chemical classes and displayed a range of physicochemical properties, exemplifying the complex chemical logic of odorant detection (Extended Data Fig. **1f,g**).

To confirm that MhORs function as ion channels, we performed whole-cell voltage-clamp recordings of HEK cells expressing MhOR5 and found that the odorant eugenol elicited slowly-activating inward currents. In outside-out patches, eugenol evoked small-conductance single channel openings that rapidly flickered between the closed and open state, resembling the activity of canonical heteromeric insect olfactory receptors^6,30^ (Fig. **1d**). Together, these data indicate that MhORs can autonomously assemble as homo-tetrameric ligand-gated ion channels that display the diverse chemical tuning profiles typical of this receptor family. We focus on MhOR5 as it is activated by a wide array of structurally diverse odorants, offering an inroad to investigate the molecular basis of promiscuous chemical recognition.

### Structure of the MhOR5 homo-tetramer

We used single-particle cryo-electron microscopy to elucidate the structure of the MhOR5 tetramer purified in detergent micelles in the absence of any ligand and obtained a density map at 3.3Å resolution (Fig. **1e,f** and Extended Data Fig. **3**–**5**). Density was well-defined, allowing us to unambiguously build a model for the majority of the protein, with the exception of several small extra-membranous loops and the short intracellular N-terminus and extracellular C-terminus (Extended Data Fig. **5c**). Despite the fact that MhOR5 shares only ∼18% amino-acid conservation with the Orco from the wasp, *Apocrypta bakeri*^30^, the structures of these two receptors display striking similarity, both in the fold of each heptahelical subunit as well as in the tetrameric organization of the subunits within the membrane plane (Extended Data Fig. **6**). As in Orco, each MhOR5 subunit contributes a single helical element (S7b) to form a central ion conduction pathway, while their S0-S6 helices form a loosely packed domain that projects radially away from the pore axis, like the spokes of a pinwheel. Within the membrane, the contacts between MhOR5 subunits are minimal and confined to the pore, whereas ∼75% of the residues that form inter-subunit interactions reside within the intracellular ‘anchor’ domain, formed from the intertwined S4-S7 helices of all four subunits (Extended Data Fig. **7**). Analogous to the Orco structure, the tightly-packed anchor domain of MhOR5 exhibited the highest local resolution (Extended Data Fig. **4c**), supporting a structural role in stabilizing the loosely assembled S0-S6 transmembrane domains within the lipid bilayer. The limited sequence conservation across neopteran ORs and Orcos maps to residues predominantly within the pore and anchor domain^30^, underscoring how the unique architecture of this receptor family can accommodate a high degree of sequence diversification while maintaining the same overall fold.

### Odorant binding leads to pore opening

To explore the structural determinants of odorant gating, we next determined a 2.9Å resolution structure of MhOR5 in complex with eugenol, the highest activity ligand identified from our screen. Three-dimensional reconstruction of the bound structure immediately yielded higher resolution, apparent from early stages of data processing, as channel features were better resolved in initial 2D class averages in the eugenol-bound versus apo structure (Extended Data Fig. **3**–**5**).

In the absence of ligand, the pore of MhOR5 displays the same distinct quadrivial architecture as the Orco homotetramer^30^, in which a single extracellular pathway opens into a large aqueous vestibule near the intracellular surface of the membrane and then diverges into four lateral conduits formed at the interfaces between subunits (Fig. **2a,b**). The branched architecture of the pore provides a route for ions to flow from the extracellular solution into the cytosol, while circumnavigating the tightly packed anchor domain. In the unbound structure of MhOR5, the S7b helices coalesce to form the narrowest portion of the ion conduction pathway. In particular, Val468 protrudes into the channel lumen generating a hydrophobic constriction of ∼5.3Å in diameter, and thus serves as a gate to impede the flow of hydrated ions through the quadrivial pore (Fig. **2a,b,d**).

**Figure 2.**
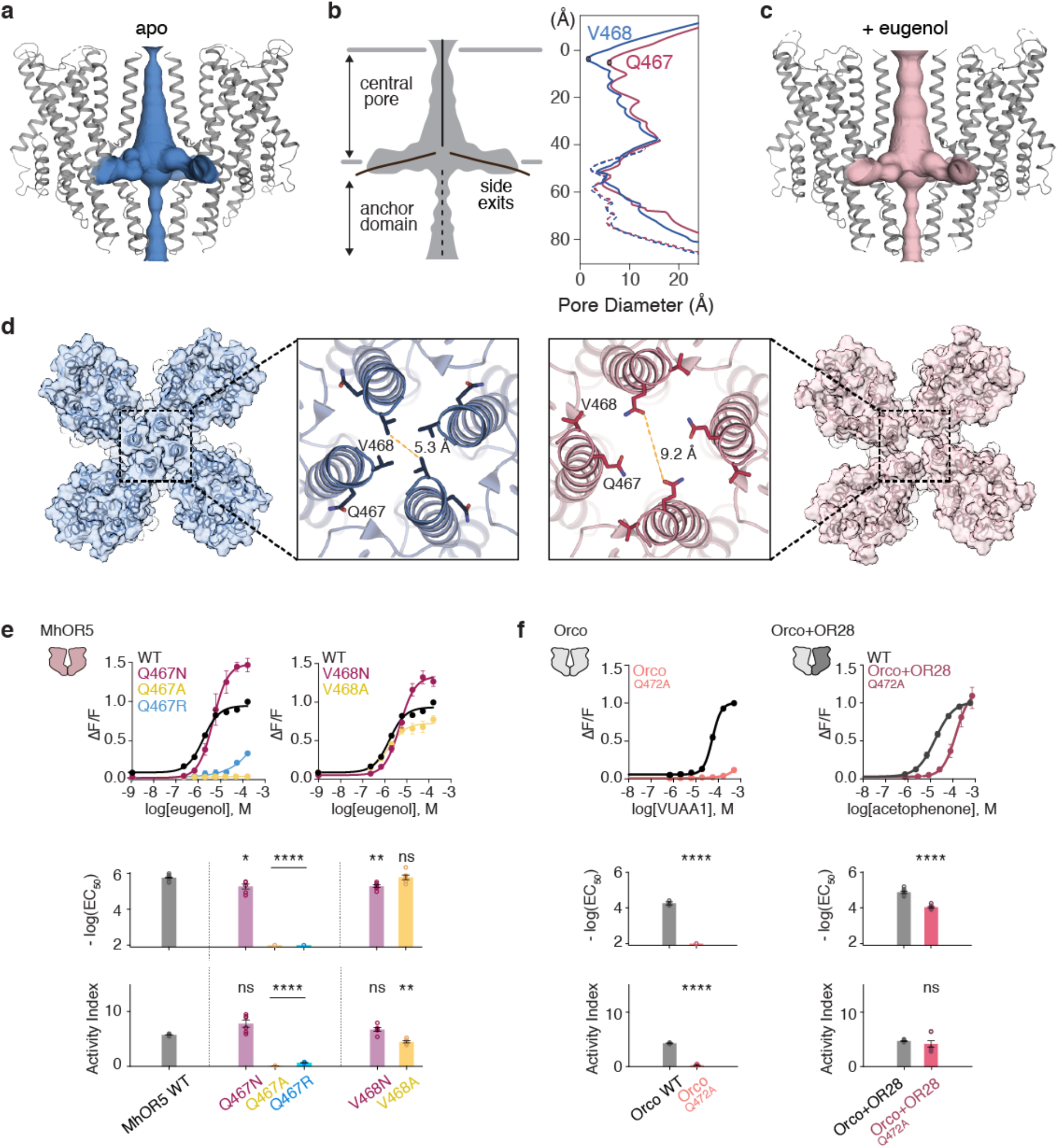
Odorant-evoked opening of MhOR5. The channel pore of MhOR5 in the unbound state (**a**, blue) and bound to eugenol (**c**, pink). **b**. The diameter of the ion conduction pathway (solid lines) and along the central 4-fold axis through the anchor domain (dashed lines). **d**. Close-up view of the pore helix S7b from the extracellular side in the apo (left, blue) and eugenol-bound (right, pink) structures, highlighting the positions of residue Gln467 and Val468. **e**. Effect of mutations of MhOR5 Val468 and Gln467. Top, dose response curves of WT and mutants. Bottom, mean log(EC_50_) and Activity Index (– log(EC_50_) ∗ max ΔF/F) with SEM for WT and mutants (n = 6-7). **f**. Pore mutants homologous to MhOR5 Gln467 in the Orco homo-tetramer or heteromeric Orco/AgOR28 complex. Top, dose response curves. Bottom, mean log(EC_50_) and Activity Index (– log(EC_50_) ∗ max ΔF/F) with SEM for WT and mutants (n = 6-7). For **e** and **f**, statistical significance was determined using one-way ANOVA followed by Dunnet’s multiple comparison tests. For mutants where the EC_50_ was incalculably high and Bartlett’s test showed non-homogenous variance, statistical significance was determined with a Brown-Forsythe test. ∗P < 0.05, ∗∗P < 0.01, ∗∗∗P < 0.001, ∗∗∗∗P < 0.0001. More information on receptor data activity in Extended Data Tables 6 and 7.

A comparison of the ion conduction pathways in the apo and eugenol-bound MhOR5 structures offers immediate insight into the conformational changes associated with gating. In the presence of eugenol, the extracellular aperture of the pore is dilated due to movement of the S7b helices away from the central pore axis (Fig. **2b-d**). The displacement of the S7b helices rotates Val468 out of the pore lumen to face the lipid bilayer, while the side chain of Gln467 rotates in to face the ion pathway. As a consequence of this small rearrangement, the chemical environment of the pore is transformed from a narrow hydrophobic constriction to a wide hydrophilic ring, 9.2Å in diameter, that can readily accommodate the passage of hydrated cations. Indeed, local resolution analysis reveals that the S7b helices display markedly increased resolution in the eugenol-bound structure in comparison to the apo structure (Extended Data Fig. **4c**), consistent with functional evidence that MhOR5 channels, like many insect olfactory receptors, display spontaneous openings even in the absence of ligand^6,16^, likely resulting in a distribution of gating states that introduces heterogeneity into the cryo-EM sample. Notably, however, the remainder of the quadrivial pore remains essentially unaltered with the addition of eugenol (Fig. **2a-c**), confirming that the tightly packed anchor domain forms a relatively stationary structural element. The dilation of the S7b helices thus appears sufficient to gate the ion conduction pathway, a small conformational change that would provide a low energetic barrier to gating, consistent with the high basal activity and low odorant affinity characteristic of most olfactory receptors^6,15,16^.

Gln467 is highly conserved across Orcos and ORs from *M. hrabei* and other basal insect species^32^ and previously identified as a component of the only identified signature sequence motif (TYhhhhhQF) diagnostic of the highly divergent insect chemosensory receptor superfamily^34,35^. Mutation of Gln467 in MhOR5 to either the smaller residue, alanine, or the positively charged residue arginine, strongly impaired function, while a more conservative mutation to asparagine resulted in enhanced activity (Fig. **2e**). In contrast, replacement of neighboring Val468 with either alanine or asparagine resulted in minimal changes to odorant activation (Fig. **2e**). In the closed structure of the Orco homotetramer^30^, the homologous residue, Gln472, points into the lipid membrane, resembling its position in the closed conformation of the MhOR5 pore. Mutation of Gln472 to alanine in Orco yielded non-functional homomeric channels (Fig. **2f**), one of the few S7b residues in Orco intolerant to such a perturbation^30^, consistent with a conserved and critical role in gating and/or ion permeation across members of this receptor family. Notably, the function of the Q472A Orco mutant could be partially rescued by co-expression with an OR from *Anopheles gambiae* (Fig. **2f**), indicating that this mutant can fold and function in the context of the heteromeric assembly and underscoring the intrinsic robustness of the Orco and OR complex where both subunits contribute to a shared ion conduction pathway^30,36^ and can compensate for variations in the pore sequence.

### Architecture of the odorant-binding site

Within the transmembrane domain of each MhOR5 subunit, the S2, S3, S4 and S6 helices splay apart to form a 15 Å-deep pocket within the extracellular leaflet of the bilayer (Fig. **3a,b**). Clearly defined density consistent with the size and shape of eugenol lies at the base of this pocket enclosed within a hydrophobic box constructed from several large aromatic and hydrophobic residues, with Trp158 forming the lid, Tyr91 and Tyr383 forming its base, and flanked by Tyr380 on one side and by Met209, Ile213 and Phe92 on the other (Fig. **3c** and Extended Data Fig. **8c**). In the apo structure, the density for some of these amino-acids was diffuse (Extended Data Fig. **8b**), which could be attributed to the overall lower resolution of this structure or to conformational flexibility when no odorant is bound. The lower resolution of the pocket in the apo state precluded us from defining the path that eugenol may take to enter the pocket or whether in the absence of an added ligand, the cavity is partially occupied, for example by a component of the buffer. Nevertheless, binding of eugenol stabilized the constellation of residues lining the pocket, leading to well-defined density and allowing for unambiguous mapping of the side chains that form the binding site.

**Figure 3.**
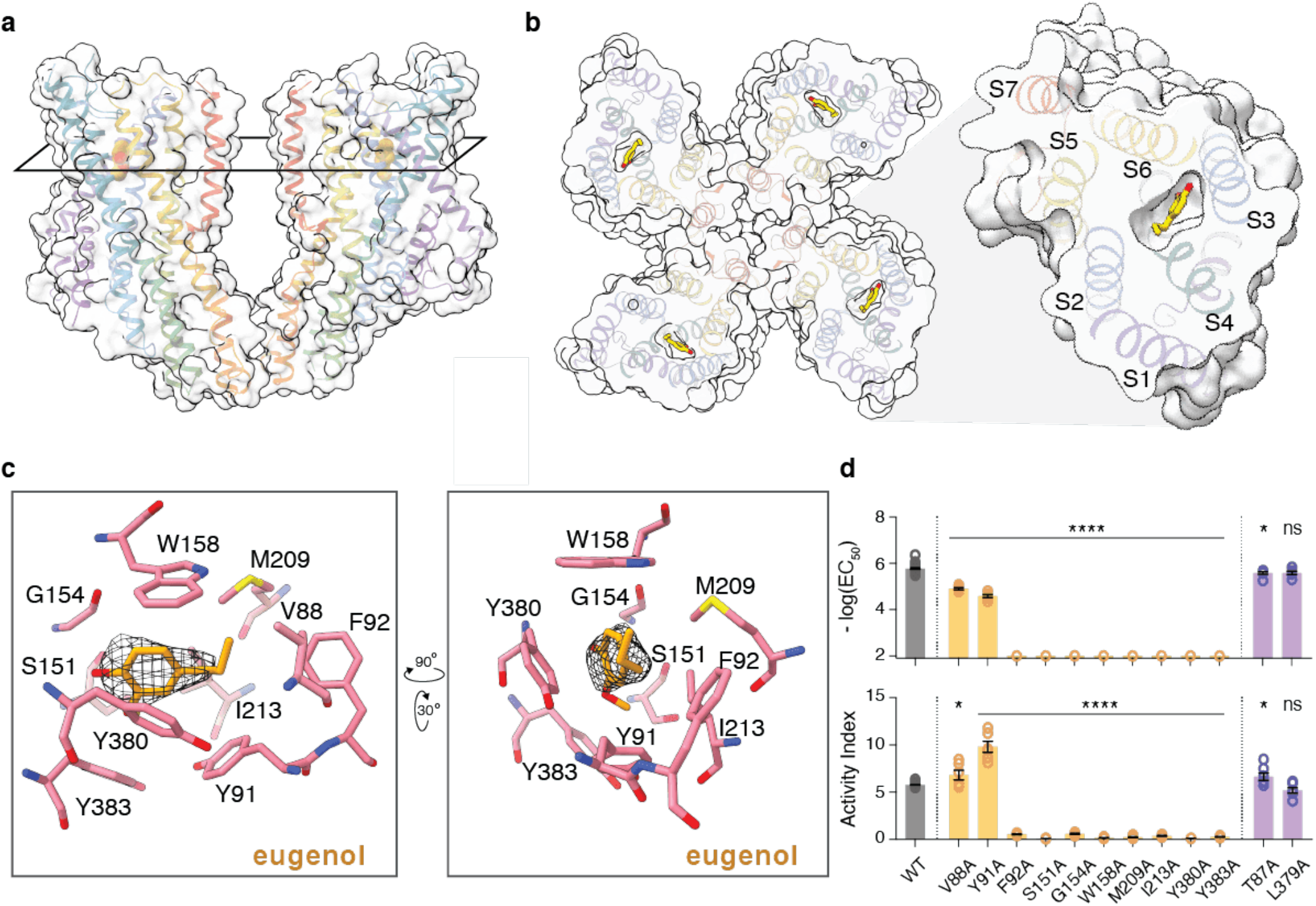
Architecture of the odorant-binding site in MhOR5. **a**. Side view of two subunits of MhOR5 with the plane through the binding pocket indicating the position of the cross section shown in (b). **b**. Top view of MhOR5 dissected at the binding pocket, with an expanded view of a single subunit on the right. Eugenol shown in stick representation within the pocket. **c**. Two views of the binding pocket. Residues in contact with eugenol are shown in pink, eugenol is shown in yellow, and cryo-EM density shown as black mesh. **d**. Mutagenesis of residues in contact with eugenol (gold) and two neighboring residues (purple) that face outward from the pocket. Mean log(EC_50_) and Activity Index shown with SEM (n = 6-7). Statistical significance was determined using one-way ANOVA followed by Dunnet’s multiple comparison tests. For mutants where the EC_50_ was incalculably high and Bartlett’s test showed non-homogenous variance, statistical significance was determined with a Brown-Forsythe test. ∗P < 0.05, ∗∗P < 0.01, ∗∗∗P < 0.001, ∗∗∗∗P < 0.0001. Dose-response curves and summary table of receptor data are available in Extended Data Fig. 10 and Extended Data Table 6, respectively.

To explore the potential binding modes of eugenol, we used computational docking methods^37^ and performed a broad grid search spanning the majority of the transmembrane domain. This analysis identified a series of closely related eugenol poses that all displayed uniformly favorable docking scores and fit well into the experimental density (Extended Data Fig. **9a**). Differentiating between these poses is challenging at this resolution given that eugenol, like most odorants, is low in molecular weight with few distinguishing structural features that can be used to orient it within the density. However, across all the top poses, eugenol was predicted to bind via comparable interactions, only these were mediated by different hydrophobic or aromatic residues within the pocket. For example, in most predicted poses, eugenol’s benzene ring was stabilized through Π-stacking interactions, but these could be alternatively mediated by Trp158, Tyr91, or Tyr380 which lie on opposing faces of the binding pocket. In every pose, eugenol also formed extensive hydrophobic interactions with a distinct but overlapping complement of aliphatic and aromatic side chains. Moreover, although eugenol’s hydroxyl was consistently oriented towards the only polar amino-acid lining the pocket (Ser151), none of the predicted poses adopted a geometry that allowed eugenol to form hydrogen bonds with any of the surrounding residues. Therefore, across these closely-related poses, eugenol recognition appears to similarly rely on non-directional hydrophobic interactions formed with a distributed array of residues lining the binding pocket. While only one of these poses may be energetically favored, past structural studies have revealed how a single odorant can bind in different orientations within the hydrophobic cavity of an odorant binding protein^38,39^, raising the possibility that eugenol may likewise sample from multiple degenerate binding modes in MhOR5.

To functionally corroborate the eugenol binding site, we identified 10 amino acids whose side chains were positioned in close proximity to the ligand density—Val88, Tyr91, Phe92, Ser151, Gly154, Trp158, Met209, Ile213, Tyr380 and Tyr383—and found that mutation of any of these residues to alanine strongly altered eugenol signaling, as indicated by a significant decrease in their apparent binding affinity (EC_50_) and/or Activity Index (Fig. **3db** Extended Data Fig. **10a**). Interestingly, the F92A, S151A, M209A, I213A and Y380A mutations also displayed increased baseline activity (Extended Data Fig. **10d**), suggesting that these residues are required to stabilize the closed conformation. Mutation of adjacent residues that project away from the binding site—Thr87 and Leu379—had minimal impact on eugenol activation (Fig.**3d**, Extended Data Fig. 1**0a**), underscoring the specificity of these perturbations to odorant-dependent gating.

How does odorant binding gate the pore? A comparison of the unbound and eugenol-bound structures indicates that in addition to the dilation of the pore, smaller conformational changes are distributed throughout the transmembrane portion of the S0-S6 helices (Extended Data Fig. **11a**). While the delocalized nature of these small rearrangements makes it challenging to delineate how odorant binding is coupled to pore opening, one potential route is through the S5 helix, which is uniquely positioned to transduce conformational changes in the binding pocket to gating of the ion pathway: S5 runs parallel to both the S7b helix that lines the pore and the S6 helix that contributes key residues to the odorant-binding pocket, namely Tyr380 and Tyr383. Upon eugenol binding, these three helices move in concert away from the central axis of the channel and towards the binding pocket, a conformational change that displaces the S7b helices outward to gate the ion conduction pathway (Extended Data Fig. **11a,b**). Close to the extracellular surface of the membrane, the S5 and S7 helices interact through a pair of hydrophobic residues, Tyr362 and Leu465, which are highly conserved as hydrophobic amino-acids across insect olfactory receptors and evolutionarily coupled^40^, pointing to a coordinated role in channel function. These hydrophobic residues remain tightly packed as the S7b helix moves into an open configuration (Extended Data Fig. **11b**), suggesting how they may contribute to coupling conformational rearrangements within the odorant binding pocket to the dilation of the pore during gating. Mutation of either Tyr362 or Leu465 to alanine impaired eugenol activation of the receptor or shifted it to higher odorant concentrations, whereas mutation of Tyr362 to phenylalanine had no effect (Extended Data Fig. **11c**), supporting a model where hydrophobic interactions at this position contribute to gating.

### Structural determinants of odorant specificity

MhOR5 is activated by a wide range of odorants (Extended Data Fig. **1f-g**), raising the question as to whether its broad tuning arises from different ligands binding to structurally distinct sites on the receptor or flexibly interacting with the same binding site. To explore this, we determined the 2.9Å structure of MhOR5 in complex with the repellent DEET, which serves as a potent and structurally distinct activator of MhOR5 and other neopteran ORs^41,42^, allowing us to investigate the diversity of binding modes used by different ligands. In the DEET-bound structure, the S7b helices of the pore were dilated to a diameter of 8.7Å (Extended Data Fig. **12**) indicating that different odorants elicit a common conformational change to open the pore. Density corresponding to DEET localized to the same binding pocket as eugenol, encased within the same box-like configuration of aromatic and aliphatic side chains (Fig. **4a-c** and Extended Data Fig. **8d**). Like eugenol, computational docking of DEET yielded multiple poses with nearly equivalent docking scores that fit the experimental density well. While each of the top poses was predicted to adopt a distinct orientation, all were stabilized through a similar complement of hydrophobic and/or Π-stacking interactions (Extended Data Fig. **9b**). Although we cannot resolve whether DEET adopts only one or multiple conformations within the binding pocket, these observations further reinforce how non-directional hydrophobic interactions may contribute to flexible chemical recognition, allowing different odorants like eugenol and DEET to bind to the same structural locus or potentially enable a single odorant to sample from multiple poses within the binding cavity.

To explore whether the broader panel of MhOR5 ligands may exploit this same chemical logic, we examined how several of their physicochemical descriptors correlated with MhOR5 activity. Using multiple regression analysis, we found that no single metric was strongly predictive of MhOR5 agonism, in accord with the notion that the molecular receptive range of a receptor reflects a complex combination of metrics^14,43,44^. However, the molecular descriptors that best accounted for MhOR5 activity were low polar surface area, low water solubility, and low potential for forming hydrogen-bonds (Extended Data Fig. **13a**), consistent with our structural observations of a geometrically simple binding site where diffuse hydrophobic interactions dominate. While hydrophobicity is a ubiquitous feature of volatile odorants^44^, MhOR1 agonism was far less correlated with these same descriptors, suggesting they play a heterogenous role in shaping the tuning of different receptors (Extended Data Fig. **13b**). Furthermore, the top 31 MhOR5 agonists identified in our panel were all predicted to favorably dock within this same binding site, relying predominantly on hydrophobic interactions (Extended Data Fig. **11c**). Diverse odorants may therefore be recognized through distributed and non-directional interactions with an overlapping subset of residues in the MhOR5 binding pocket, providing a potential basis for this receptor’s broad tuning.

How does the MhOR5 binding pocket accommodate such diverse ligands? A comparison of the eugenol and DEET-bound structures reveals that the constellation of amino acids lining the binding pocket retains the same overall geometry, leaving the architecture of the hydrophobic box largely unchanged. However, a small displacement of the S4 helix results in an expansion of the pocket, likely to accommodate the longer aliphatic moiety of DEET and avoid a steric clash with the aliphatic side chain Met209 (Fig. **4c** and Extended Data Fig. **9a,b**). Functional data support these structural observations. Mutation of Met209 to smaller hydrophobic amino-acids, valine or alanine, enhanced DEET affinity, as indicated by a shift in the dose-response curve to lower concentrations of ligand (Fig. **4d** and Extended Data Fig. **10b**). The same mutations attenuated eugenol sensitivity, suggesting that this smaller odorant less optimally occupies the binding pocket in the absence of the bulky methionine side chain. Conversely, mutation of Ile213, another aliphatic S4 residue that lies in close proximity to the DEET density, to the larger residue methionine abolished DEET sensitivity but had marginal impact on eugenol signaling (Fig. **4d** and Extended Data Fig. **10c**). Structure-guided mutagenesis therefore predictably altered MhOR5’s sensitivity to these two odorants.

**Figure 4.**
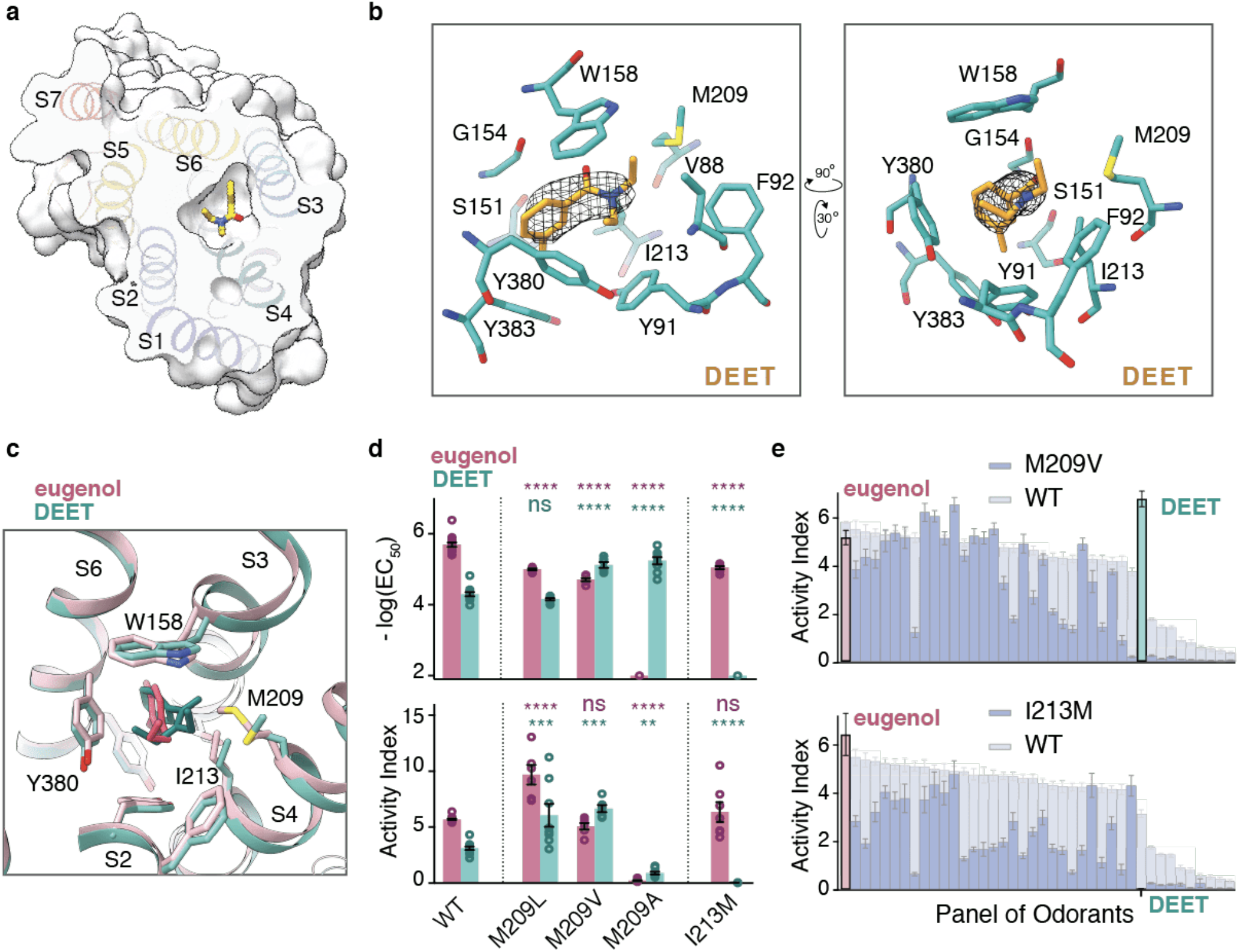
Structure-based mutagenesis retunes MhOR5. **a**. Cross-section of the binding pocket of a MhOR5 subunit in complex with DEET. DEET is shown in stick representation within the pocket. **b**. Two views of the binding pocket (same orientations as shown in Fig. 3c). DEET is shown in yellow with cryo-EM density shown as black mesh. **c**. Overlay of the MhOR5 binding pockets of DEET-bound (teal) and eugenol-bound (pink), with both ligands shown. **d**. Effect of mutating Met209 and Ile213 into residues of variable side-chain length on eugenol and DEET signaling. Mean log(EC_50_) and Activity Index ± SEM (n = 6-7). Statistical significance was determined using one-way ANOVAs followed by Dunnet’s multiple comparison tests comparing mutants to their respective wild-type controls for each ligand. For mutants where the EC_50_ was incalculably high and Bartlett’s test showed non-homogenous variance, statistical significance was determined with a Brown-Forsythe test. ∗P < 0.05, ∗∗P < 0.01, ∗∗∗P < 0.001, ∗∗∗∗P < 0.0001. **e**. Tuning curves of M209V (top) and I213M (bottom). The tuning curve of wild-type MhOR5 in response to a panel of 40 odorants is shown in grey, sorted from maximum to minimum Activity Index, and the M209V and I213M mutant tuning curves are shown overlaid in dark grey (n = 5-6 for mutants and n = 10-17 for wild-type with each ligand) with eugenol (pink) and DEET (teal) highlighted. Dose-response curves are available in Extended Data Fig. 10, and summary table of receptor data can be found in Extended Data Table 6,8, and 9.

Given that our analyses together point to a shared structural locus for odorant recognition in MhOR5, we examined the effects of the I213M and M209V mutations using a larger panel of 40 odorants. We found that each point mutation extensively altered receptor tuning, with some odorants exhibiting impaired signaling, while other odorants displayed enhanced sensitivity as a consequence of the same mutation (Fig. **4e**). Changes in odorant tuning for either mutant did not adhere to a simple obvious logic, consistent with the complexity of physicochemical properties that define MhOR5 agonism (Extended Data Fig. **13**) and reinforcing that both the global geometry and local chemical environment of the binding pocket contribute to tuning a receptor’s chemical sensitivity. Subtle modifications of the MhOR5 binding pocket therefore can broadly reconfigure the receptor’s chemical tuning, supporting a model in which different odorants are recognized by a common binding site.

## Discussion

The broad tuning of olfactory receptors is central to the detection and discrimination of the vast chemical world. Here we offer structural insight into how such flexible chemical recognition is achieved. We find that MhOR5 detects a wide array of odorants through a single, promiscuous binding site that recognizes the overall physicochemical properties of each odorant, rather than being tuned to any of its specific chemical or molecular features. Our work therefore supports that receptor tuning depends on the stereochemistry of its ligands^18,45^, yet does not adhere to the classic lock-and-key mechanism that governs many receptor-ligand interactions.

Several structural features of the MhOR5 binding pocket endow the receptor with its broad chemical tuning. First, odorant binding relies predominantly on hydrophobic interactions, which lack the strict geometric constraints inherent to other intermolecular associations, like hydrogen bonds, that frequently mediate ligand recognition. Second, the distributed arrangement of hydrophobic and aromatic residues across multiple surfaces of the binding pocket further relaxes orientational constraints by allowing odorants to form comparable interactions with many of its faces. Third, the S2, S3, and S6 helices are rich in aromatic residues that confer structural rigidity to the pocket, enabling it to retain its overall shape in the presence of structurally diverse ligands like DEET and eugenol. Thus, the simple geometry of the binding site alleviates the requirement for shape complementarity, imposing minimal restriction on the shape of odorants that can bind. Finally, a degree of structural flexibility is nevertheless provided by aliphatic residues clustered on the S4 helix that can rearrange to accommodate larger ligands like DEET. Indeed, mutation of Met209 and Ile213 in S4 to different-sized hydrophobic residues resulted in predictable and opposing changes to eugenol and DEET sensitivity, consistent with their role in defining the dimensions of the binding cavity. The prevalence of these comparatively weak intermolecular interactions is compatible with the relatively low affinity of most odorants^15,16,46^. By capturing multiple gating states of MhOR5, we show that dilation of the S7b helices by only a few angstroms is sufficient to gate the ion conduction pathway. This subtle conformational change would present a low energetic barrier to opening, consistent with the high baseline activity of insect olfactory receptors and their brief flickering transitions between the closed and open states^6,16^.

Computational docking analyses support our structural observations, reinforcing how the non-directional nature of hydrophobic interactions and distinct geometry of the binding pocket can accommodate a wide array of structurally and chemically diverse ligands. The flexibility we observe is reminiscent of structural studies of odorant binding proteins (OBPs) that ferry diverse hydrophobic ligands through the sensory neuron lymph, and similarly rely on distributed hydrophobic interactions for their promiscuous chemical tuning^39^. Crystallographic studies of OBPs reveal that the same odorant can adopt alternative conformations within its binding pocket^38,47^, supporting a model in which odorants may sample from different binding modes within a hydrophobic cavity. Indeed, in engineered ligand-binding pockets that rely on hydrophobic interactions for molecular recognition, the same ligand has been shown to bind in multiple conformations^48^, further emphasizing how the directionality of ligand binding is poorly constrained in the absence of hydrogen-bonding or electrostatic interactions.

Promiscuous ligand binding is a hallmark of many chemosensory receptors^1,3^. Although the structure of a mammalian olfactory receptor has yet to be elucidated, odorant recognition has been proposed to rely on distributed hydrophobic and non-directional interactions within a deep transmembrane pocket^19,49,50^, mirroring how flexible chemical detection is mediated in MhOR5. Thus, structurally and mechanistically distinct olfactory receptor families appear to rely on similar principles for their broad chemical tuning, pointing to common constraints in how diverse hydrophobic molecules are recognized. Additional mechanisms for odorant recognition certainly exist, in particular for receptors that are selectively tuned to ethologically relevant chemical classes^51–53^, including more hydrophilic compounds likes acids or amines^54,55^, or cues whose perceptual meaning is singular and invariant, like pheromones^52^. Whether stricter odorant specificity relies on distinct intermolecular binding modes, variations in the geometry of the binding pocket, or both, remains to be elucidated.

A key observation from our work is that multiple odorants are recognized within the same binding site in MhOR5. Residues implicated in dictating odorant specificity in distantly related neopteran insect ORs map to MhOR5’s binding pocket^56–61^ (Extended Data Fig. **14**), indicating that it represents a conserved and canonical locus for odorant detection across this highly divergent family. Unexpectedly, we find that the insect repellent DEET binds to this same site, offering structural corroboration that ORs represent one of DEET’s molecular targets. Natural variation in DEET sensitivity has previously identified to a single residue^57^ which maps to the MhOR5 binding pocket, lending support to the proposal that DEET may exploit the promiscuity of diverse ORs and serve as a molecular ‘confusant’ by scrambling the olfactory code^57,62^. Other modulators of olfactory receptors, like VUAA1 which inhibits MhOR5, cannot favorably dock within its binding pocket due to its much larger size, suggesting that insect olfactory receptors may harbor additional points of allosteric modulation that expand their signaling mechanisms.

Several important implications arise from our observation that diverse odorants bind to the same locus within a receptor and are recognized by shared structural determinants. First, even single conservative mutations within the binding site can broadly reconfigure the receptor’s chemical tuning, likely facilitating the rapid evolution of receptors with distinct ligand specificity^2,56,61,63^. However, this also poses a significant evolutionary constraint as individual binding-site mutations will likely have a pleotropic effect on the representation of multiple odorants, potentially serving to broadly reconfigure the odor code. Second, at the level of neural coding, our observations underscore the fundamental challenge of evolving receptor repertoires that are tuned with sufficient breadth to support detection of the immense chemical world but also conferred with sufficient specificity to mediate odor discrimination. The promiscuity and arbitrary nature of odorant recognition at the receptor level likely imposes significant selective pressures on the structure and function of olfactory circuits, driving the evolution of synaptic and circuit mechanisms to decorrelate, decode, and impose meaning onto combinatorial patterns of receptor activity^64,65^. Odor discrimination is thus transformed from a biochemical problem at the receptor level to a neural coding problem within the brain.

Finally, our work provides insight into the evolution of the insect olfactory system. Although the role of MhORs in chemosensory detection in the jumping bristletail remains unknown, we demonstrate that they can autonomously assemble and function as homomeric odorant-gated ion channels. The striking structural similarity of MhOR5 with Orco underscores how the architecture of this family is maintained despite extensive sequence diversification. These observations support the proposal that MhORs lie at the ancestral origin of the insect olfactory receptor family, from which the massive and divergent family of Orco/OR heteromeric channels arose^31,32^. Why neopteran ORs became obligate heteromers with Orco remains enigmatic but presumably reflects the fact that Orco confers structural stability to the complex, thereby relaxing evolutionary constraints on the ORs and allowing them to further diversify, ultimately supporting the flexible detection and discrimination of an enormous and changing chemical world.

## Methods

### Expression and purification of MhOR5

The coding sequence of *M. hrabei* OR5 (MhOR5) was synthesized as a gene fragment (Twist). Residues Lys2 to Pro474 were cloned into a pEG BacMam vector^66^ containing superfolder GFP^67^, an N-terminal Strep II tag, and an HRV 3C protease site for cleavage. SF9 cells (ATCC CRL-1711) were used to produce baculovirus containing the MhOR5 construct, and the virus, after three rounds of amplification, was used to infect HEK293S GnTI^-^ cells (ATCC CRL-3022)^66^. HEK293S GnTI^-^ cells were grown at 37°C with 8% carbon dioxide in Freestyle 293 medium (Gibco) supplemented with 2% (v/v) fetal bovine serum (Gibco). Cells were grown to 3 × 10^6^ cells/mL and infected at a multiplicity of infection of ∼1. After 8-12 h, 10 mM sodium butyrate (Sigma-Aldrich) was added to the cells and the temperature was dropped from 37°C to 30°C for the remainder of the infection duration. 72 h after initial infection, cells were harvested by centrifugation, washed with phosphate-buffered saline (pH 7.5; Gibco), weighed and flash frozen in liquid nitrogen. Pellets were stored at −80°C until thawed for purification.

For purification, cell pellets were thawed on ice and resuspended in 20 mL of lysis buffer per gram of cells. Lysis buffer was composed of 50 mM HEPES/NaOH (pH 7.5), 375 mM NaCl (together, 4X HBS), 1 μg/mL leupeptin, 1 μg/mL aprotinin, 1 μg/mL pepstatin A, 1 mM phenylmethylsulfonyl fluoride (PMSF; all from Sigma-Aldrich) and ∼3 mg DNase I (Roche). MhOR5 was extracted using 0.5% (w/v) n-dodecyl β-D-maltoside (DDM; Anatrace) with 0.1% (w/v) cholesterol hemisuccinate (CHS; Sigma-Aldrich) for 2 h at 4°C. The mixture was clarified by centrifugation at 90,000 x g and the supernatant was added to 0.5 mL StrepTactin Sepharose resin (GE Healthcare) per gram of cells and rotated at 4°C for 2 h. The resin was collected and washed with 10 column volumes (CV) of 1X HBS with 0.025% (w/v) DDM and 0.005% (w/v) CHS (together, SEC buffer). MhOR5 was eluted by adding 2.5 mM desthiobiotin (DTB) and cleaved overnight at 4°C with HRV 3C Protease (EMD Millipore). Sample was concentrated to ∼5 mg/mL and injected onto a Superose 6 Increase column (GE Healthcare) equilibrated in SEC buffer. Peak fractions containing MhOR5 were concentrated until the absorbance at 280 nm reached 5-6 (approximately 5 mg/mL) and immediately used for grid preparation and data acquisition. For the eugenol bound structure, peak fractions were pooled, and eugenol (Sigma Aldrich, CAS#97-53-0) dissolved in dimethylsulfoxide (DMSO; both Sigma-Aldrich) was added for a final odor concentration of 0.5 mM, and the complex was incubated at 4°C for 1 h. The maximum DMSO concentration was kept below 0.07%. The complex was then concentrated to approximately 5 mg/ml and used for grid preparation. For the DEET bound structure, sample from the overnight cleavage step was concentrated to ∼5 mg/ml and injected into the Superose 6 Increase column equilibrated in SEC buffer with 1 mM DEET (Sigma Aldrich, CAS#134-62-3). Peak fractions were concentrated to ∼5 mg/ml and used immediately for grid preparation.

### Cryo-EM sample preparation and data acquisition

Cryo-EM grids were frozen using a Vitrobot Mark IV (FEI) as follows: 3 μl of the concentrated sample was applied to a glow-discharged Quantifoil R1.2/1.3 holey carbon 400 mesh gold grid, blotted for 3-4 s in > 90% humidity at room temperature, and plunge frozen in liquid ethane cooled by liquid nitrogen.

Cryo-EM data were recorded on a Titan Krios (FEI) operated at 300 kV, equipped with a Gatan K2 Summit camera. SerialEM^68^ was used for automated data collection. Movies were collected at a nominal magnification of 29,000X in super-resolution mode resulting in a calibrated pixel size of 0.51 Å/pixel, with a defocus range of approximately –1.0 to –3.0 μm. Fifty frames were recorded over 10 s of exposure at a dose rate of 1.22 electrons per Å^2^ per frame.

Movie frames were aligned and binned over 2 × 2 pixels using MotionCor2^69^ implemented in Relion 3.0^70^, and the contrast transfer function parameters for each motion-corrected image were estimated using CTFFIND4^71^.

#### Apo structure

2 datasets were collected with 4,050 micrographs in dataset A and 3,748 micrographs in dataset B. Processing was done independently for each dataset in the following way: particles were picked using a 3D template generated in an initial model from a data set of 5,000 particles picked in manual mode. A total of 562,794 (dataset A) and 536,145 (dataset B) particles were subjected to 2D classification using RELION-3.0^70^. 210,833 (dataset A) and 183,061 (dataset B) particles from the best 2D classes were selected and subjected to 3D classification imposing C4 symmetry and adding a soft mask to exclude the detergent micelle after 25 iterations. One class from each dataset containing 44,884 (dataset A) and 43,788 (dataset B) particles was clearly superior in completeness and definition of the transmembrane domains. These particles were subjected to 3D refinement with C4 symmetry, followed by Bayesian polishing and CTF refinement. The polished particles from both datasets were exported to cryoSPARC v2^72^ and processing continued with the joined dataset of 88,672 particles. In cryoSPARC, further heterogenous refinement resulted in a single class with 49,832 particles that were subjected to particle subtraction with a micelle mask. Non-uniformed refinement of subtracted particles imposing C4 symmetry yielded the final map with an overall resolution of 3.3Å estimated with a cutoff for the Fourier Shell Correlation of 0.143^73^.

#### Ligand-bound structures

Processing for the eugenol-bound and DEET-bound structures occurred through the following pipeline: 4,410 (eugenol) and 4,365 (DEET) micrographs were collected and used to pick 461,254 (eugenol) and 787,448 (DEET) particles that were extracted unbinned and exported into cryoSPARC v2. In cryoSPARC, several rounds of 2D classification resulted in 221,339 (eugenol) and 180,874 (DEET) particles that were used to generate an initial model with 4 classes with no imposed symmetry. These models were inputted as templates of a heterogenous refinement with no imposed symmetry, from which one (eugenol) and two (DEET) final classes were selected containing 129,031 (eugenol) and 121,441 (DEET) particles. These particles were refined and exported to RELION 3.0 where they were subjected to a round of 3D classification with no imposed symmetry. The best class from this 3D classification contained 54,900 (eugenol) and 56,191 (DEET) particles that were subjected to Bayesian polishing and CTF refinement. Polished particles were then imported into cryoSPARC v2 and subjected to particle subtraction. Final non-uniform refinement with C4 symmetry imposed resulted in the final maps with overall resolution of 2.9Å in both cases, estimated with a cutoff for the Fourier Shell Correlation of 0.143. In all cases, the four-fold symmetry of the channel was evident from the initial 2D classes without having imposed symmetry and refinements without imposed symmetry produced 4-fold symmetric maps.

### Model Building

The Cryo-EM structure of Orco (pdb 6c70) was used as a template for homology modeling of MhOR5 using Modeller^74^, followed by manual building in Coot^75^. The 3.3Å density map of the apo was of sufficient quality to build the majority of the protein, with the exception of the S3-S4 and S4-S5 loops, the 13 N-terminal residues and the 5 C-terminal residues. The models were refined using real space refinement implemented in PHENIX^76^ for 5 macro-cycles with fourfold non-crystallographic symmetry applied and secondary structure restraints applied. The eugenol- and DEET-bound models were refined including the ligands, that were placed as starting point within the corresponding density in a pose that was obtained through docking methods (described below). Model statistics were obtained using MolProbity. Models were validated by randomly displacing the atoms in the original model by 0.5Å, and refining the resulting model against half maps and full map^77^. Model to map correlations were determined using phenix.mtriage. Images of the maps were created using UCSF ChimeraX^78^. Images of the model were created using PyMOL^79^ and UCSF ChimeraX^78^.

### Docking analysis

All compounds were docked using Glide^37,80^ implemented in Maestro (Schrödinger, suite 2020). Briefly, the model was imported into Maestro and prepared for docking. A 20Å^3^ cubical grid search was built centered in the region of observed ligand density. Ligand structures were imported into Maestro by their SMILES unique identifiers and prepared using Epik^81^ to generate their possible tautomeric and ionization states, all optimized at pH 7.0 ± 2. All ligands were docked within the built grid, and the top poses that best fit the density are presented in Extended Data Fig. 8. The top activators scored with values between −7.4 and −4. While all activators docked with negative scores, some non-activators also docked with favorable scores. For example, caffeine docked favorably despite the molecule not activating the channel in our functional experiments. As docking does not incorporate dynamics of the receptor, it is not expected that docking will correlate homogeneously or monotonically with experimentally determined activity of ligands. At most a qualitative agreement can be expected.

### Structure analysis

Residues at subunit interfaces were identified using PyMOL as any residue within 5Å of a neighboring subunit (Extended Data Fig. 7). The pore diameters along the central axis and lateral conduits were calculated using the program HOLE^82^ (Fig. 2b, Extended Data Fig. 12a). Two calculations were performed for each structure: one along the central four-fold axis (central pore) and another between subunits near the cytosolic membrane interface (lateral conduits). The plots in Fig. 2b and Extended Data Fig. 12a show the diameter along the central axis of the main conduit and the lateral conduit.

### Electrophysiology

The full length MhOR5 sequence was cloned into a pEG BacMam vector. HEK293 cells were maintained in high-glucose DMEM supplemented with 10% (v/v) fetal bovine serum (FBS) and 1% (v/v) GlutaMAX (all Gibco) at 37°C with 5% (v/v) carbon dioxide. Cells were plated on 35 mm tissue culture-treated Petri dishes 72-48 h before recording, and infected with GFP-tagged MhOR5 24-48 h before recording. Electrodes were drawn from borosilicate patch glass (Sutter Instruments) and polished (MF-83, Narishige Co.) to a resistance of 3-6 MΩ when filled with pipette solution. Analog signals were filtered at 2 kHz using the built-in 4-pole Bessel filter of a Multiclamp 700B patch-clamp amplifier (Molecular Devices) in patch mode and digitized at 20 kHz (Digidata 1440A, Molecular Devices). Signals were further filtered offline at 1 kHz for analysis and representations.

Whole-cell and single-channel recordings in **Error! Reference source not found**.d were performed using an extracellular (bath) solution composed of 135 mM NaCl, 5 mM KCl, 2 mM MgCl2, 2 mM CaCl2, 10 mM glucose, 10 mM HEPES-Na/HCl (pH 7.3, 310 mOsm/kg) and an intracellular (pipet) solution composed of 150 mM KCl, 10 mM NaCl, 1 mM EDTA-Na, 10 mM HEPES-Na/HCl (pH 7.45, 310 mOsm/kg). Single-channel recordings were done in excised outside-out mode. Eugenol was dissolved in DMSO at 150 mM and diluted to 0.1 mM in the extracellular solution. Solutions were locally perfused using a microperfusion system (ALA Scientific Instruments).

### Cell-based GCaMP fluorescence calcium flux assay

All DNA constructs used in this assay were cloned into a modified pME18s vector with no fluorescent marker. Each transfection condition contained 0.5 ug of a plasmid encoding GCaMP6s (Addgene #40753) and 1.5 ug of the plasmid encoding the appropriate olfactory receptor, diluted in 250 μL OptiMEM (Gibco). In experiments with heteromeric olfactory receptors, the total amount of DNA was 1.5 ug, in a ratio of 1:1 of Orco/OR. These were diluted in a solution containing 7 μL Lipofectamine 2000 (Invitrogen) and 250 μL OptiMem, followed by a 20 min incubation at room temperature. HEK293 cells were maintained in high-glucose DMEM supplemented with 10% (v/v) fetal bovine serum (FBS) and 1% (v/v) GlutaMAX at 37°C with 5% (v/v) carbon dioxide. Cells were detached using trypsin and resuspended to a final concentration of 1 × 10^6^ cells/mL. Cells were added to each transfection condition, mixed and added to 2 × 16 wells in a 384-well plate (Grenier CELLSTAR). 4-6 h later, a 16-port vacuum manifold on low vacuum was used to remove the transfection medium, replaced by fresh FluoroBrite DMEM (Gibco) supplemented with 10% (v/v) FBS and 1% (v/v) GlutaMAX. 24 h later, this medium was replaced with 20 μL of reading buffer (20 mM HEPES/NaOH (pH 7.4), 1X HBSS (Gibco), 3 mM Na_2_CO_3_, 1 mM MgSO_4_, and 2 or 5 mM CaCl_2_) in each well. The calcium concentration was optimized for each receptor to account for their differences in baseline activity: for experiments with MhOR5 and MhOR5 mutants, reading buffer contained 2 mM CaCl_2_, while 5 mM CaCl_2_ was used for MhOR1, Orco and Orco/AgOR28 heteromers. The fluorescence emission at 527 nm, with excitation at 480 nm, was continuously read by a Hamamatsu FDSS plate reader. After 30 s of a baseline recording, an optimized amount of odorant solution— 10 μL for all MhOR-containing experiments or 20 μL for all Orco-containing experiments—was added to the cells and read for 2 min. All solutions were warmed to 37°C before beginning.

Seven ligand concentrations were used for each transfection condition in sequential dilutions of 3, alongside a control well of only reading buffer. Ligands were dissolved in DMSO to 150 mM, then diluted with reading buffer to a highest final-well concentration of 0.5 mM (DMSO never exceeded 0.5%). Water soluble ligands (arabinose, caffeine, denatonium, glucose, MSG, sucrose) were dissolved directly into reading buffer. If experimental data indicated a more sensitive response than this range, the concentration was adjusted accordingly. Ligand concentrations for mutants were the same as corresponding wild-type OR. Each plate contained a negative control of GCaMP6s transfected alone and exposed to eugenol for MhOR5 and VUAA1 for Orco experiments. Additionally, each plate included the corresponding wild-type OR with its cognate ligand–MhOR5 and MhOR1 with eugenol, Orco with VUAA1, and Orco/AgOR28 with acetophenone–as a positive control to account for plate-to-plate variation in transfection efficiency and cell count. A control of DMSO alone was also tested to ensure no activity effects were due to the solvent. Each concentration of ligand was applied to four wells of a transfection condition, which were averaged and considered a single biological replicate.

The baseline fluorescence (F) was calculated as the average fluorescence of the 30 s before odor was added to the plate. Within each well, ΔF was calculated as the difference between the average of the last 10 seconds of fluorescence and the baseline F. ΔF/F was then calculated as the ΔF divided by the baseline fluorescence (F). Finally, the ΔF/F for each concentration was normalized to the maximum ΔF/F value of the corresponding positive control present on each plate: MhOR5 and MhOR1 with eugenol, Orco with VUAA1, and Orco/AgOR28 with acetophenone to account for inevitable variations in transfection efficiency and cell counts across different plates. The normalized ΔF/F averaged across all experiments for a given condition is the value used to construct the dose-response curves in all plots and Fig. s (Fig. s 1c, 2e,f, Extended Data Fig. s 2d, 10a-c, 12c). All wild-type curves come from the same plates as the experimental data in the same plot.

For all experiments, GraphPad Prism 8 was used to fit the dose-responses curves to the Hill equation from which the EC_50_ of the curve was extracted. Three metrics were used to characterize the dose-response curve for each ligand: Activity Index, log(EC_50_) and max ΔF/F. EC_50_ is the concentration of ligand at which the response reaches the midpoint. For conditions where EC_50_ was too high for the dose-response curve to reach saturation and therefore could not be fitted to a Hill equation, a value of −2 was assigned to the EC_50_, which is over an order of magnitude higher than the highest concentration employed. Max ΔF/F is the maximum response achieved at the highest concentration. Activity Index is defined as the negative product of log(EC_50_) and max ΔF/F, as follows:

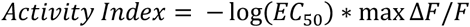

### Gels and small-scale transfections

For Western blots and FSEC traces (Extended Data Fig. 1b,c, 10d), HEK293 cells were maintained in high-glucose DMEM supplemented with 10% (v/v) fetal bovine serum (FBS) and 1% (v/v) GlutaMAX at 37°C with 5% (v/v) carbon dioxide. Cells were detached using trypsin and plated in 6-well plates at a concentration of 0.4 × 10^6^. 24 h later, cells were transfected with 3 ug of DNA in the same superfolder GFP-containing pEG BacMam vector used for large-scale purification and 9 μL FuGene HD (VWR) diluted in 150 μL OptiMEM and added dropwise to the cells after a 5 min incubation. 24 h later, cells were checked for GFP fluorescence, rinsed with phosphate-buffered saline, and harvested by centrifugation. Cells were either frozen at - 20°C or used immediately.

Cell pellets were thawed briefly and resuspended in 200 μL of lysis buffer containing 4X HBS, an EDTA-free protease inhibitor cocktail (Roche), and 1 mM PMSF. The protein was extracted for 2 h at 4°C by adding 0.5% (w/v) DDM with 0.1% (w/v) CHS after 10 s sonication in a water bath. This mixture was then clarified by centrifugation and filtered. The supernatant was added to a Shimadzu autosampler connected to a Superose 6 Increase column equilibrated in SEC buffer. An aliquot of the supernatant was also used to run SDS-PAGE (Bio-Rad, 12% Mini-PROTEAN TGX) and Blue Native(BN)-PAGE (Invitrogen, 3-12% Bis-Tris) gels. Gels were transferred using Trans-Blot Turbo Transfer Pack (Bio-Rad) and blocked overnight. The following day, gels were stained with rabbit anti-GFP polyclonal antibody (Life Technologies; 1:20,000), washed, incubated with anti-rabbit secondary antibody (1:10,000), and imaged with ImageLab.

### Lifetime sparseness calculation

The lifetime sparseness^83,84^ measure was used to quantify olfactory receptor tuning breadth and calculated in the following way:

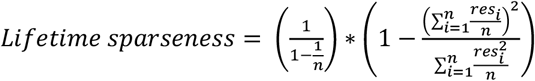

where *n* is the number of ligands in the set, *res*_*i*_ is the receptor’s response to a given ligand *i*. All inhibitory responses (values below 0) were set to 0 before the calculation^83,84^. The *D. melanogaster* OR data set come from the DoOR database^85^.

### Multiple regression analysis

A set of 11 molecular descriptors were compiled for all 54 ligands tested from PubChem, Sigma-Aldrich, ChemSpider, EPA, and The Good Scents Company, the values used are in Extended Data Table 10. A multiple regression analysis using the scikit-learn Linear Regression module was used to assess the accuracy with which the receptor activity could be predicted by individual descriptors (1-dimensional analysis) or combinations of two descriptors (2-dimensional analysis) (Extended Data Fig. 13). Due to the absence of reported metrics for some ligands – acetic acid, citric acid, MSG, sucrose, denatonium, and VUAA1 – the analysis was performed on the remaining 48 ligands. For the 1-dimensional analysis, a single variable linear regression was performed for each descriptor independently. The analysis sought to fit a linear model with coefficients w1, … wn+1 where n is the dimension of the input data. The optimal coefficient set was determined by method of residual sum of squares optimization between the observed Activity Index targets and those predicted by linear approximation using solved coefficients. This process was repeated for the 2-dimensional case, using every unique permutation of descriptors across the 11-dimensional space. As a means of assessing the predictive power of a given combination, the R^2^-value, reflecting the square of the correlation coefficient between observed and modeled values of the Activity Index, was calculated for each linear model and reported in Extended Data Fig. 13. This allowed ranking of descriptor sets based on accuracy of prediction.

### Sequence Alignments

In Fig. 1b, 28 sequences of Orcos, 82 sequences from 4 neopteran species (*Nasonia vitripennis, Drosophila melanogaster, Anopheles gambiae* and *Pediculus humanus*) and the 5 sequences of *Machilis hrabei* were aligned based on secondary structure prediction using Promals3D^86^ with minimal manual adjustment based off of the structure of MhOR5. The unrooted tree in Fig. 1b was calculated and plotted using iTOL^87^. For Extended Data Fig. 1a, the alignment between the sequences of MhOR1 and MhOR5 was done using MAFFT implemented in JalView^88^ with minimal manual adjustment based off of the structure of MhOR5. For Extended Data Fig. 6a, the sequences alignment between Abak Orco and MhOR5 was done by aligning the published structure of Abak Orco (pdb 6c70) and the structure of MhOR5 in PyMOL. All sequence alignments were visualized and plotted using JalView^88^.

## Acknowledgements

We thank R. Axel, R. MacKinnon, J. Chen, S. Klinge, B. Noro, J. Butterwick, M. Maldonado, C. McBride and members of the Ruta Lab for helpful discussion and comments on the manuscript, R. Hite, T. Walz, and J. Butterwick for advice on Cryo-EM processing, P. Stock for advice on the multiple regression analysis and P. Brand for advice on receptor alignments. We are grateful to M. Ebrahim, J. Sotiris, and H. Ng at The Rockefeller University Evelyn Gruss Lipper Cryo-Electron Microscopy Resource Center for assistance with microscope operation, and L. Ramos-Espiritu at The Rockefeller University High Throughput & Spectroscopy Resource Center for support on functional assays. This work was supported by a Leon Levy Postdoctoral Fellowship (to J.d.M) and the National Institutes of Health (R01AI103171 to V.R.).

## Author contributions

V.R. and J.d.M. conceived the study, designed experiments and wrote the manuscript, with input from M.A.Y. J.d.M. purified MhOR5, performed electrophysiology, collected and analyzed Cryo-EM data, built and refined the models, and performed molecular docking. M.A.Y. performed molecular biology, cell culture and calcium imaging assays.

## Competing interests

The authors declare no competing interests.

---

**Extended Data Fig. 1.**
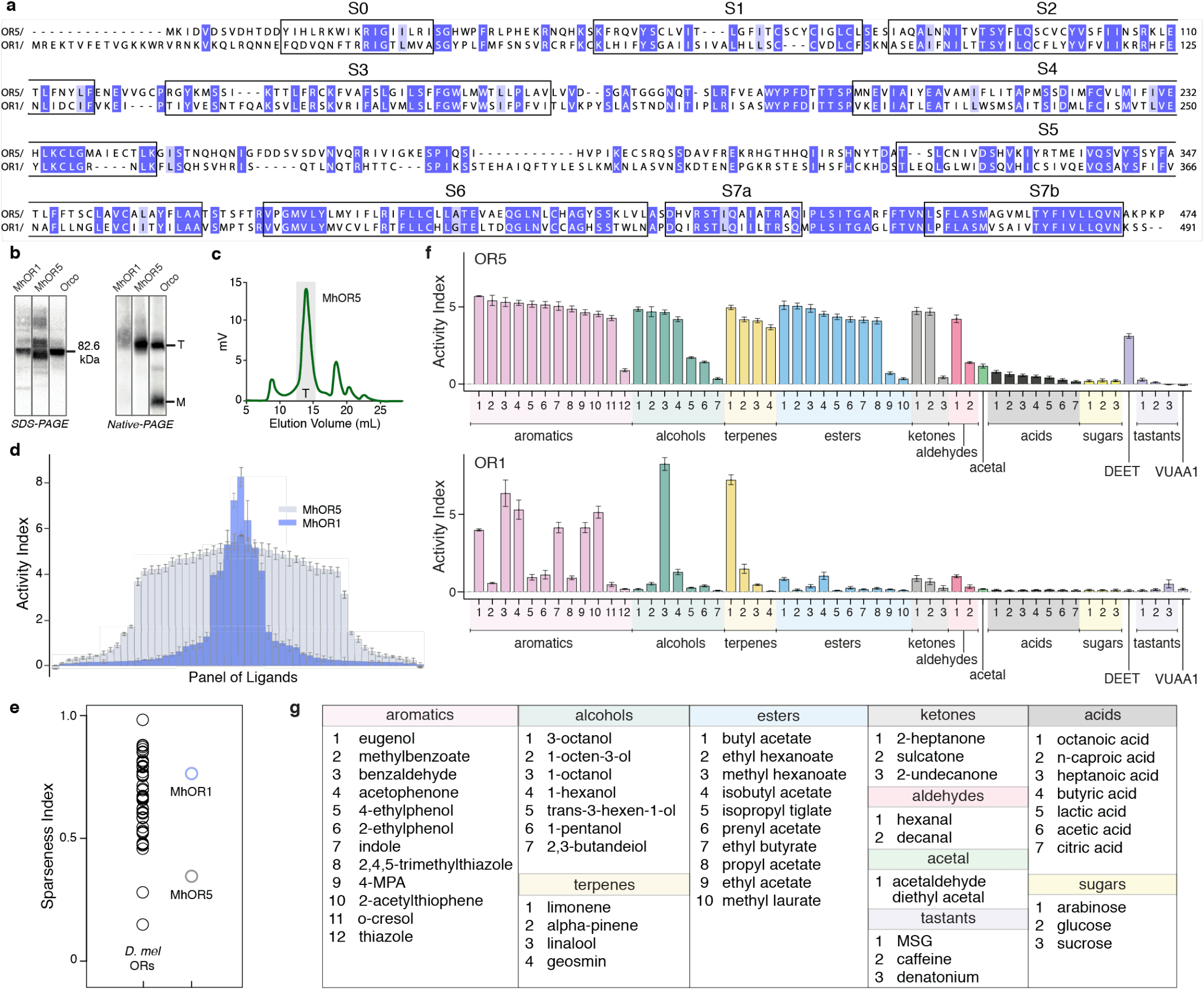
Biochemical and functional comparison of MhOR5 and MhOR1. **a**. Alignment of MhOR1 and MhOR5 protein sequences with identical (dark purple) and similar residues (light purple) highlighted. Boxes mark helical segments from MhOR5 structure. **b**. Western blots of MhOR1, MhOR5, and *A. bakeri* Orco fused to GFP and stained with anti-GFP antibodies. Left, a denaturing gel; right, a non-denaturing Blue Native gel. Position of the Orco tetramer (T) and monomer (M). **c**. Size-exclusion chromatography trace of MhOR5 with position corresponding to the tetramer stoichiometry labeled (‘T’). **d**. Receptor activity curves shown superimposed for MhOR5 (grey) and MhOR1 (blue). The same panel of 54 small molecules was used with 3 < n < 122 (median n = 11.5) for MhOR5 and 3 < n < 21 (median n = 4) for MhOR1. In both curves, Activity Index was measured as – log(EC_50_) ∗ max ΔF/F, and eugenol response was used to normalize max ΔF/F. The order of the ligands is different for each curve to highlight the breadth of tuning of each receptor. **e**. Comparison of the breadth of tuning, measured as lifetime sparseness (methods) of *D. melanogaster* ORs and *M. hrabei* ORs. Lifetime sparseness close to 0 suggests broad tuning while 1 would suggest tuning to a single ligand in the panel. **f**. Receptor activity for MhOR5 (top) and MhOR1 (bottom) with ligands (named in **g**) sorted by chemical classes. Summary tables of receptor data is available in Extended Data Table 2 (MhOR5) and 4 (MhOR1), and lifetime sparseness values can be found in Extended Data Table 5.

**Extended Data Fig. 2.**
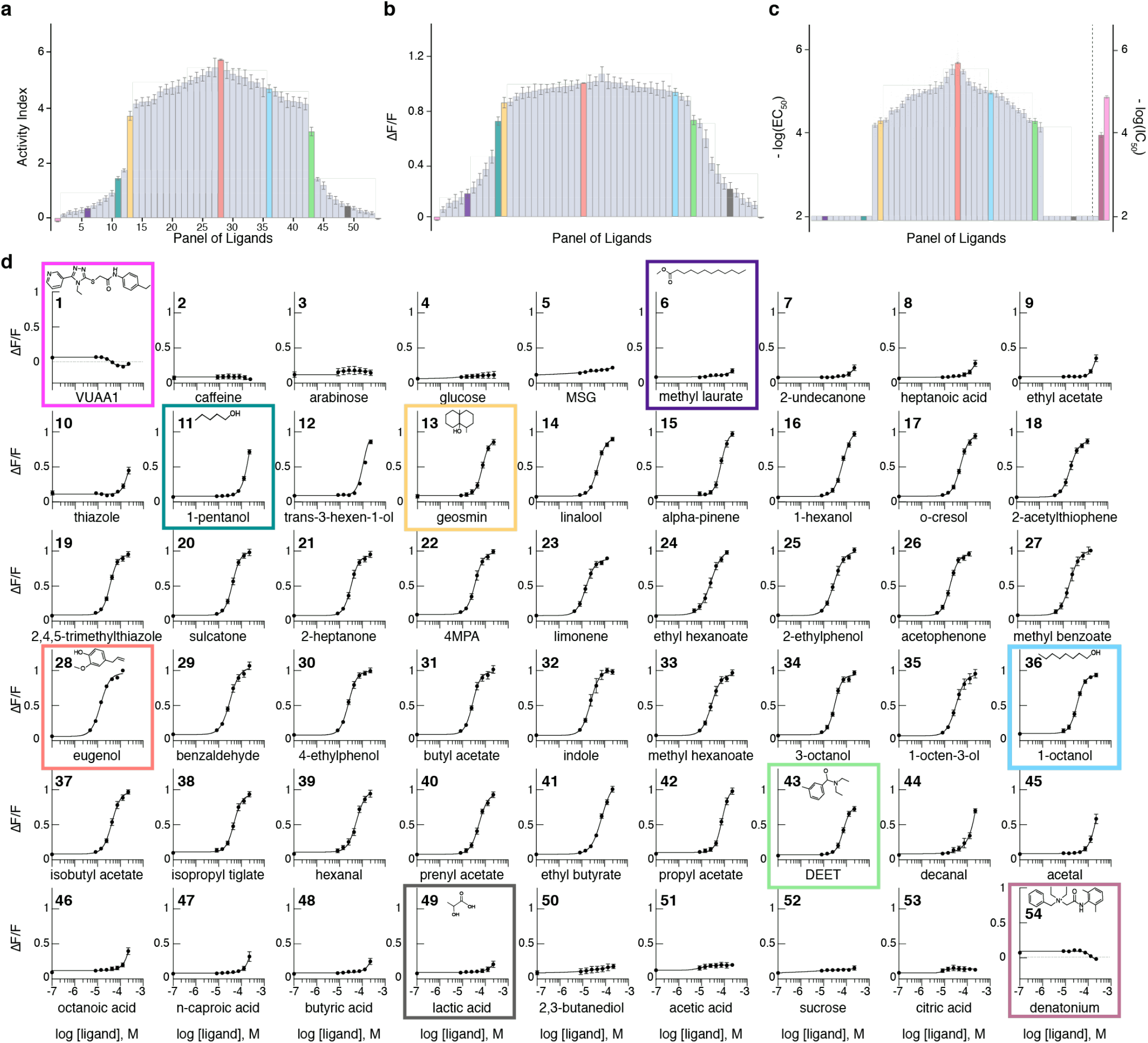
MhOR5 is activated by a broad set of odorants. Tuning curves of MhOR5 ordered by Activity Index (**a**), max ΔF/F (**b**), and – log(EC_50_) (**c**). Activity Index was calculated as − log(EC_50_) ∗ max ΔF/F. ΔF/F for each ligand is normalized to the eugenol value obtained in the same experiment to control for variation in cell count and transfection efficiency. Dose-response curves that did not saturate according to the fitted Hill equation were assigned a log(EC_50_) value of −2. In **c**, the inhibitory ligands VUAA1 and denatonium are shown as IC50. **d**. Dose-response curves for all the individual ligands shown in **a, b, c**, averaged across all experiments and shown with SEM. Numbering corresponds to the numbering of the tuning curve shown in **a**. Note that the Activity Index, which combines – log(EC_50_) and max ΔF/F captures agonism by ligands with low affinity such as 1-pentanol (teal) or sub-maximal efficacy such as DEET (green). Summary table of receptor data is available in Extended Data Table 2 and ordering of ligands in **a, b, c** found in Extended Data Table 3.

**Extended Data Fig. 3.**
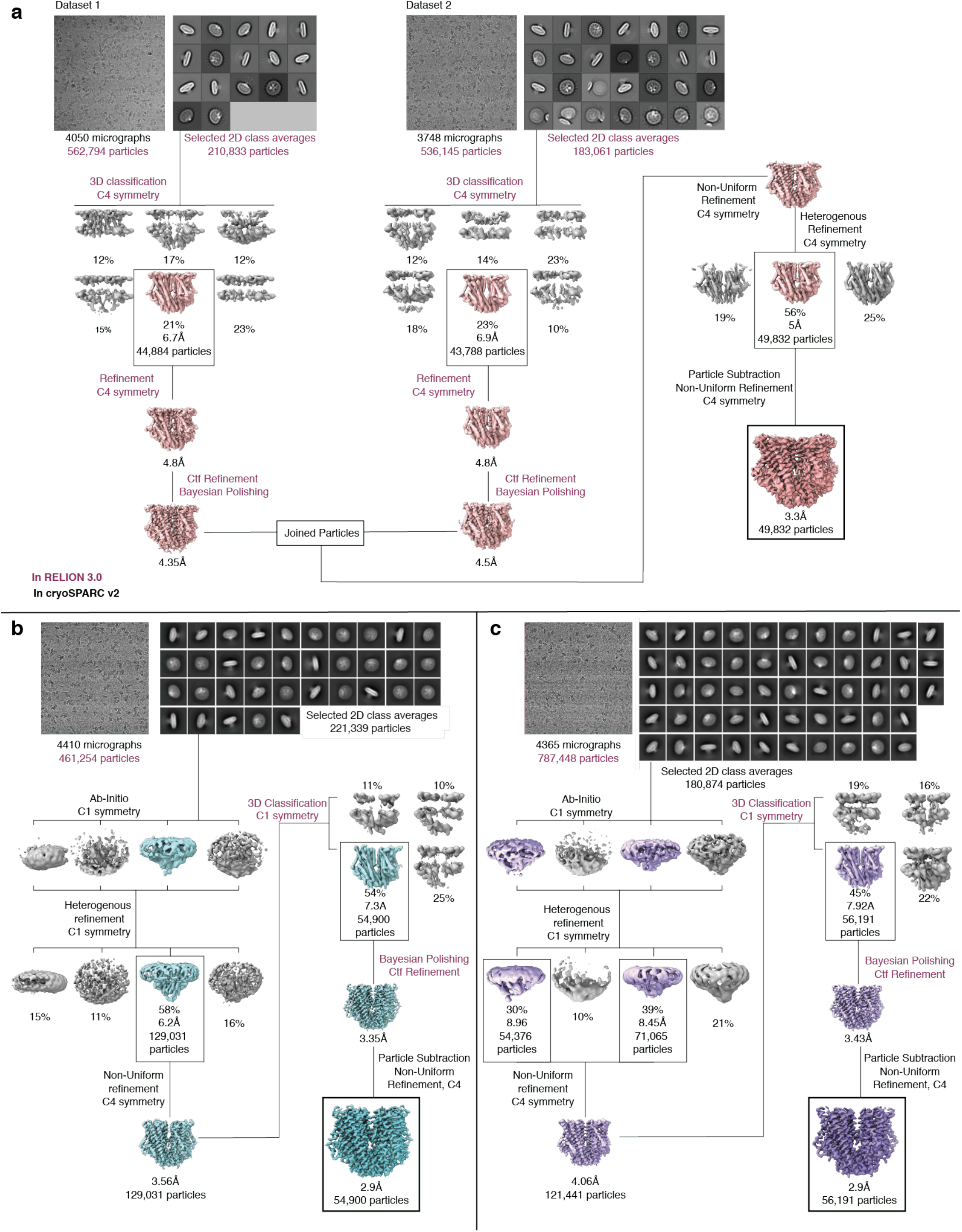
Cryo-EM data analysis. Processing pipeline for the apo structure (**a**), the eugenol-bound structure (**b**), and the DEET-bound structure (**c**).

**Extended Data Fig. 4.**
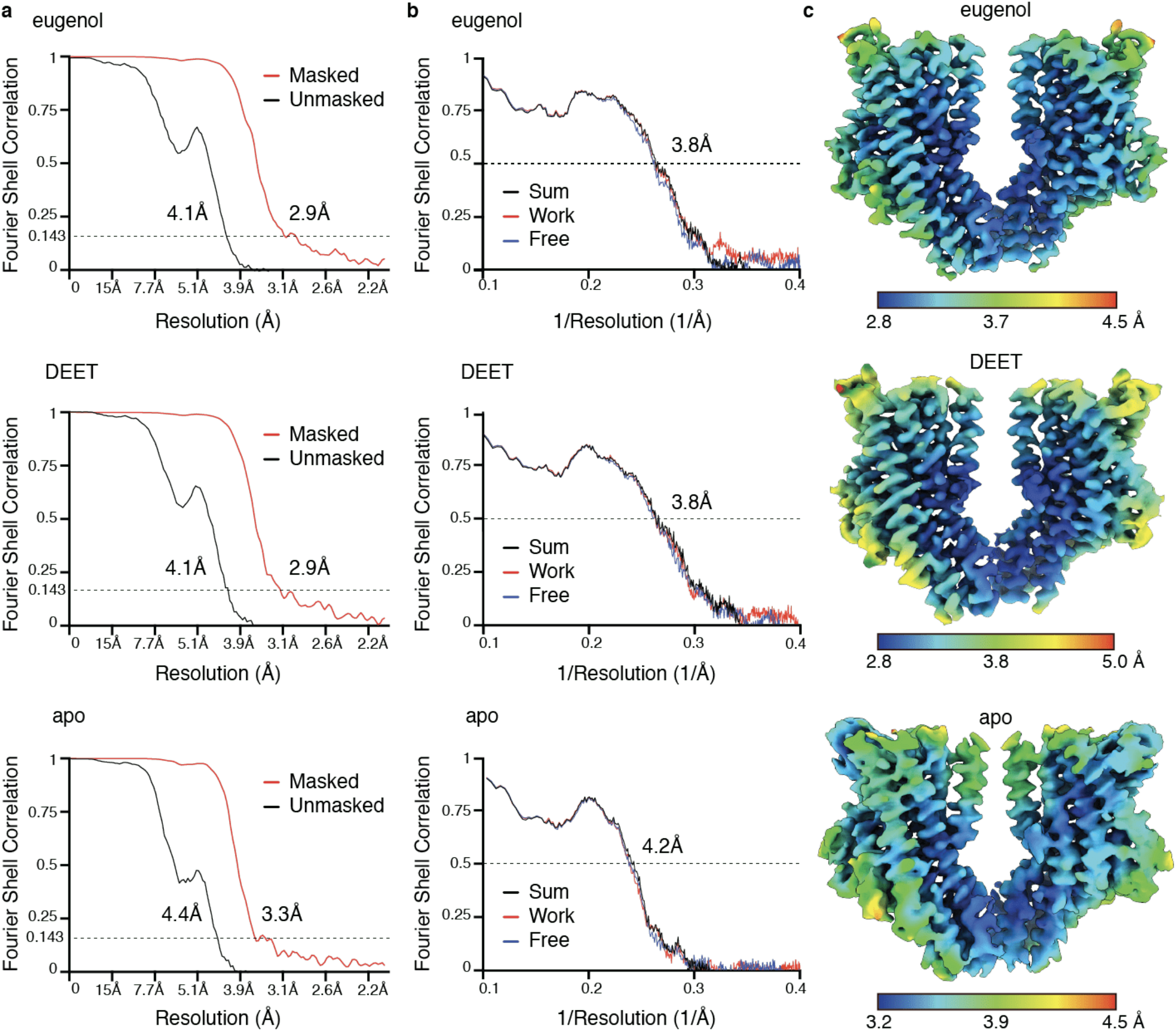
Map and model analysis and validation. **a**. Fourier shell correlation (FSC) curves for the final cryo-EM density maps, obtained with cryoSPARC v2. The horizontal dashed line intersects at 0.143, the cutoff value. **b**. FSC relationships between final map versus model (black, sum), half-map 1 versus model (red, work), and half-map 2 versus model (blue, free), calculated in phenix.mtriage. **c**. Local resolution estimation for each final map, calculated in cryoSPARC v2. Side views shown with front and back subunits removed for visualization.

**Extended Data Fig. 5.**
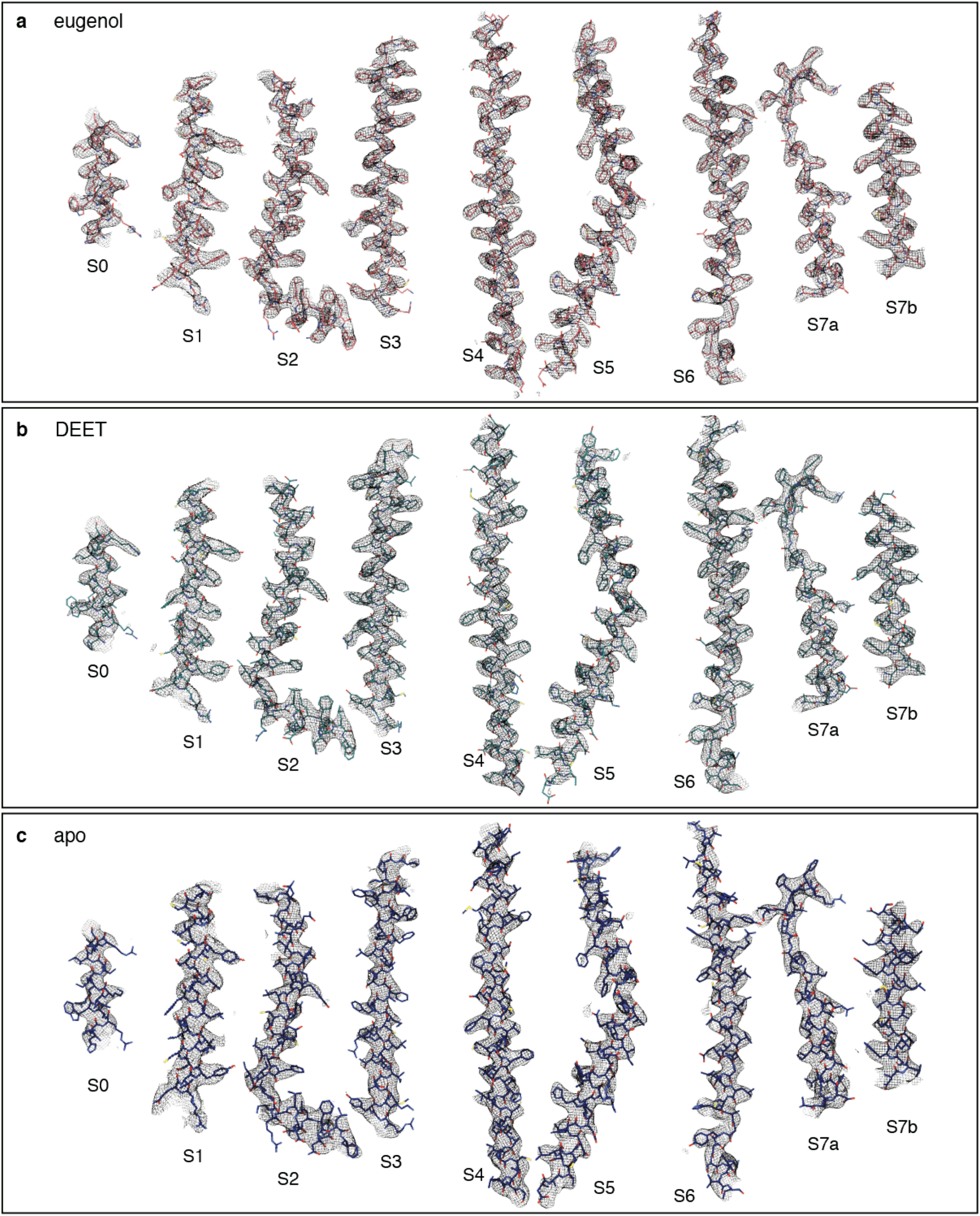
Cryo-EM density. Cryo-EM density for the modeled regions of the eugenol-bound structure (**a**), the DEET-bound structure (**b**), and the apo structure (**c**). Models are shown in stick representation within the density, with the helices denoted underneath from the N-term (S0) to the C-term (S7b).

**Extended Data Fig. 6.**
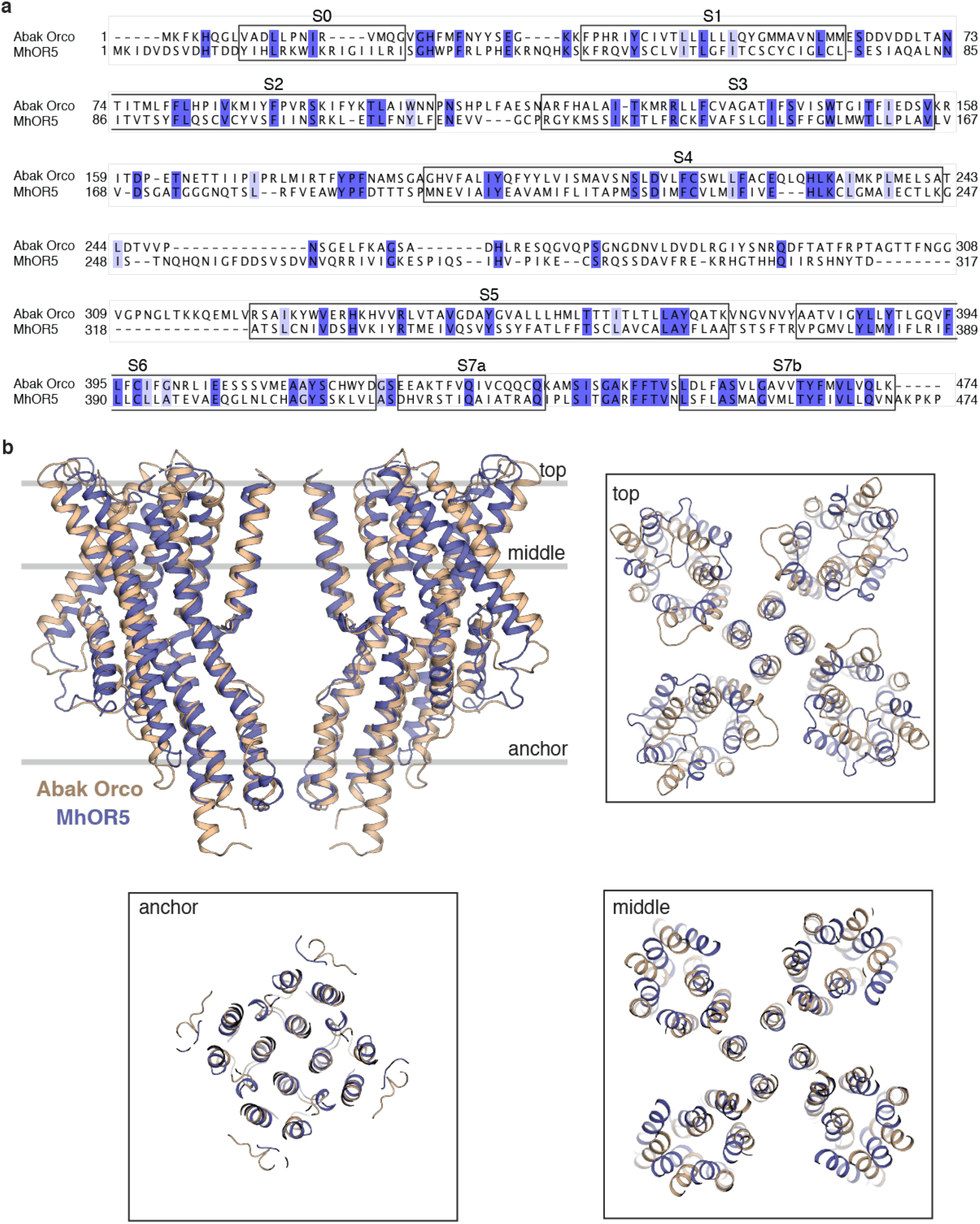
Conserved architecture of insect olfactory receptors. **a**. Sequence alignment of MhOR5 and *A. bakeri* Orco. Sequence identity (dark purple) and similarity (light purple) are highlighted. The positions of the helices in MhOR5 are marked. **b**. Structural overlay of Abak Orco (gold) and the apo state of MhOR5 (blue) from the side view (left) with grey bars indicating the positions at which cross-sections are taken for inset shown from top views (right).

**Extended Data Fig. 7.**
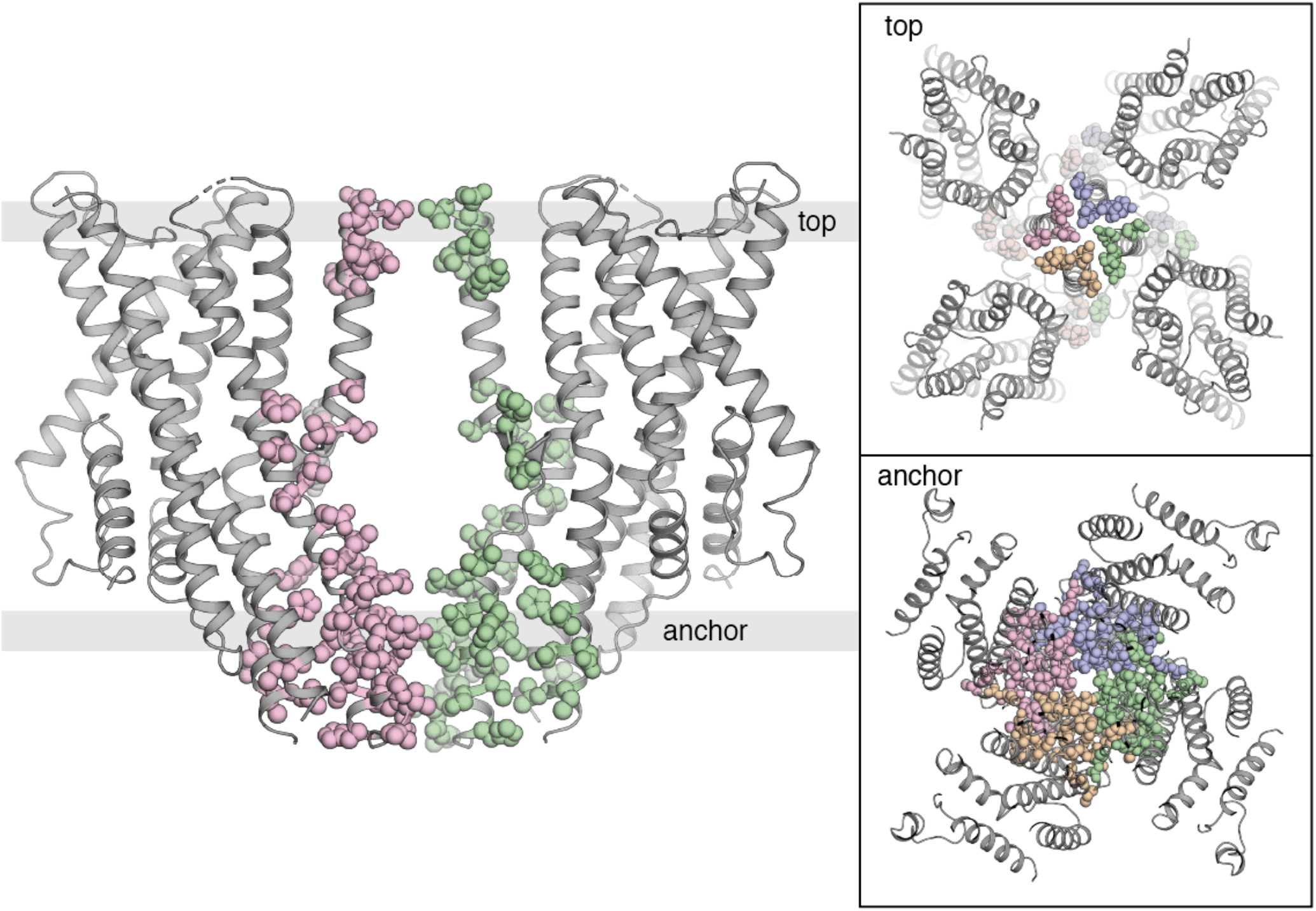
Inter-subunit interactions are concentrated in the anchor region. Side view of MhOR5 with front and back subunits removed for visualization. Residues within 5Å of residues in neighboring subunits are shows as spheres, colored by subunit. Insets (right) show top views of cross-sections taken at the top and anchor position as indicated by a grey bar in the side view (left).

**Extended Data Fig. 8.**
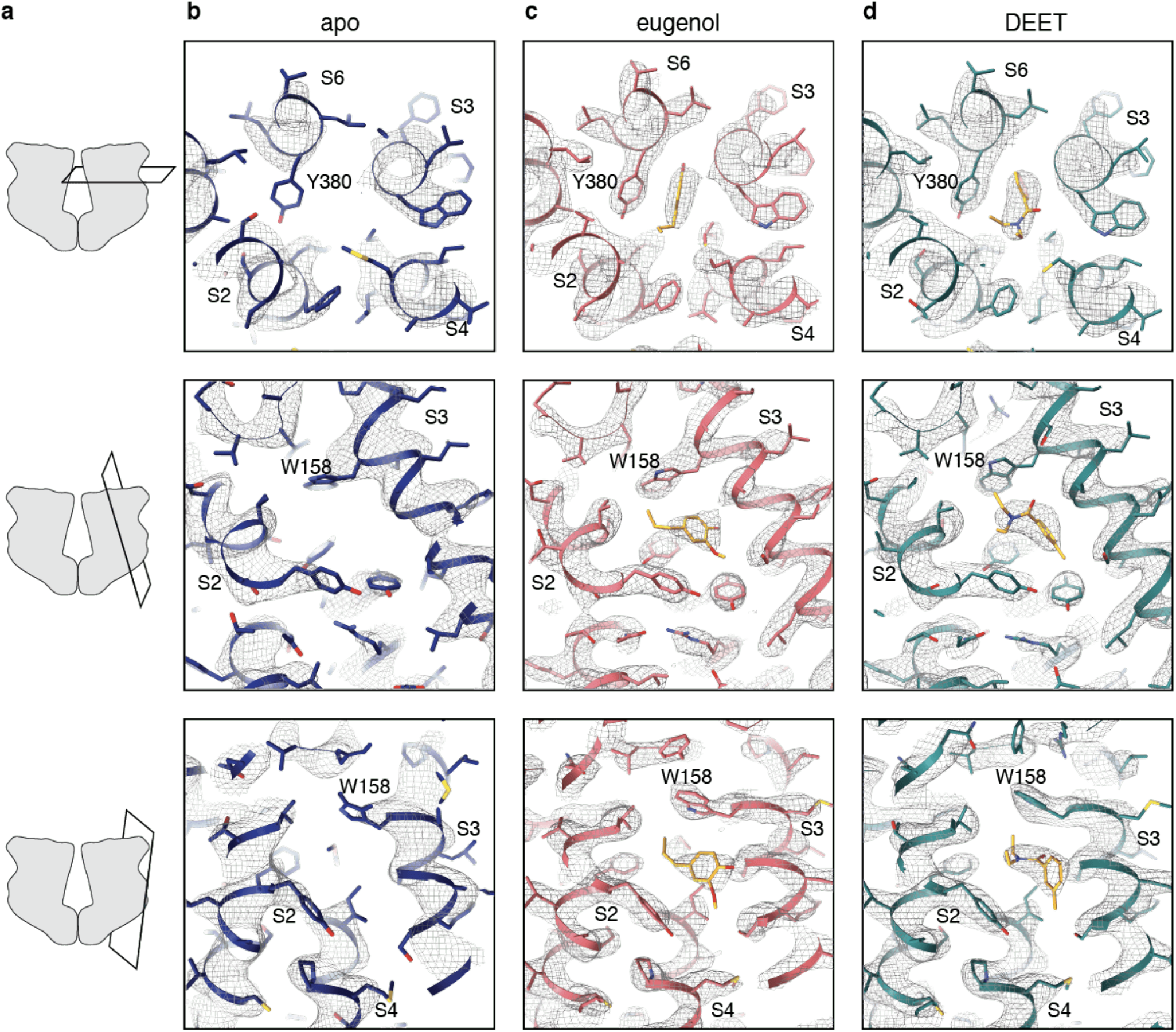
Cryo-EM density in the odorant-binding region. **a**. Schematic of the position of the three different views of the binding pocket shown in **b, c** and **d**. Model shown as ribbon and density shown as black mesh of the odorant-binding region of the apo (**b**), eugenol-bound (**c**) and DEET-bound (**d**)structures. Cryo-EM density contoured at 9 σ in all images.

**Extended Data Fig. 9.**
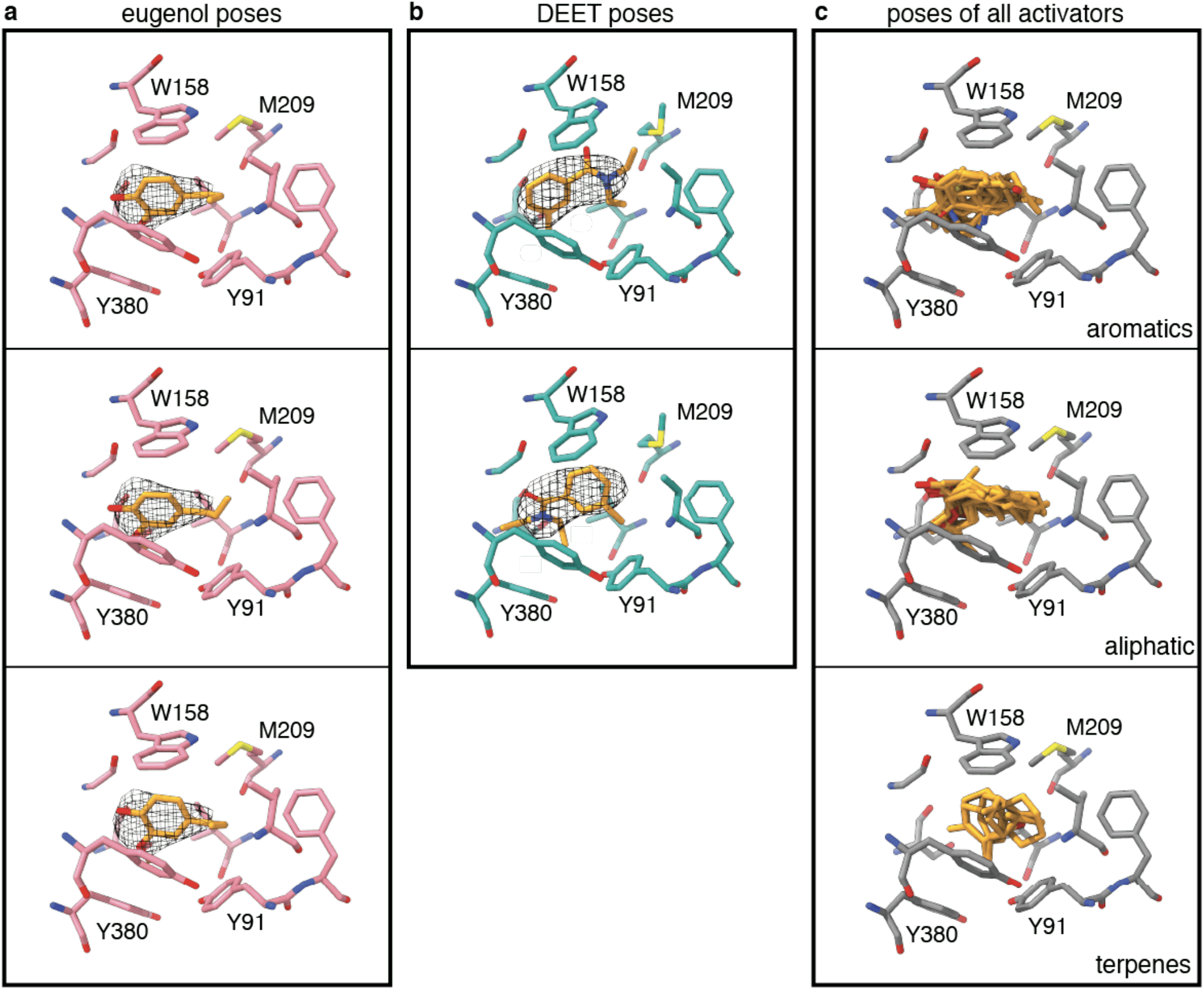
Docking of MhOR5 agonists in the binding pocket. View of the binding pocket in the eugenol bound (**a**, pink) and DEET-bound (**b**, teal) with the representative top poses for eugenol (**a**) and DEET (**b**). Cryo-EM density for eugenol (a) and DEET (b) shown as black mesh. The docking scores calculated using Glide of the eugenol poses shown in **a** are from top to bottom: −6,59, −6.71 and −6,81. The docking scores calculated using Glide of the DEET poses shown in **b** are −7.33 (top) and −7.37 (bottom). **c**, top pose for each of the agonists of MhOR5 from Glide, overlaid according to chemical class.

**Extended Data Fig. 10.**
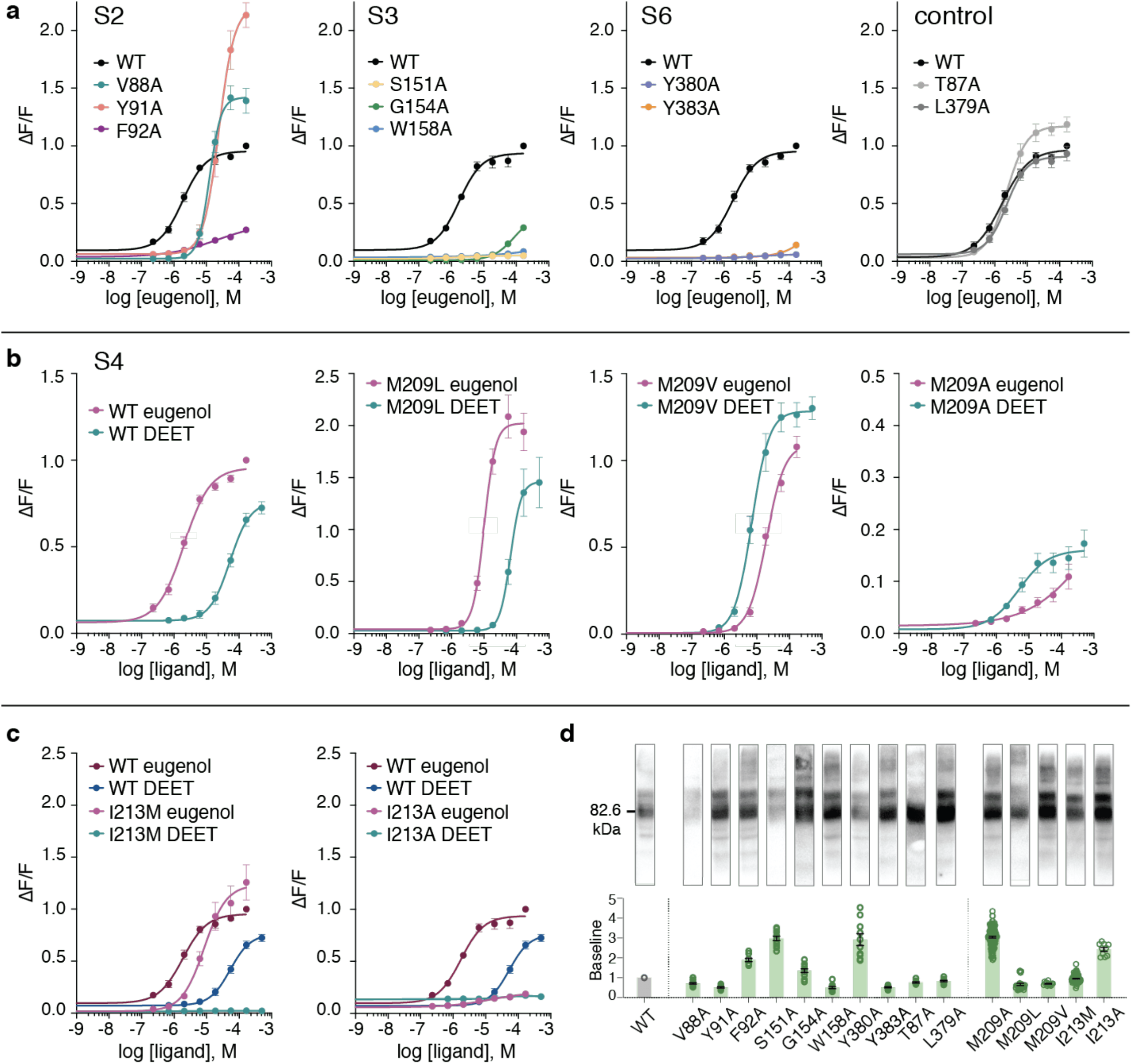
Effect of binding pocket mutations on odorant signaling, baseline activity, and protein expression. **a**. Dose-response curves of MhOR5 mutants in S2, S3, S6 helices and two control residues that neighbor the binding pocket but face away from it (n = 6-7). Each WT curve represents the corresponding controls from the same experiments. **b**. Dose-response curves of WT or M209A mutant receptor in response to eugenol (pink) and DEET (teal). **c**. Dose-response curves of WT and I213A (left) or I213M (right) mutant receptor in response to eugenol (pink) and DEET (teal). **d**. For each mutant shown in **a, b, c**, a denaturing gel showing protein expression (top) and baseline fluorescence normalized to wild-type MhOR5 (bottom). Baseline represents the mean of the first 30 s of fluorescence before the addition of any ligand. Mean and SEM displayed, with 8 < n < 116 (median n = 12). Summary of receptor data is available in Extended Data Tables 6 (eugenol) and 8 (DEET).

**Extended Data Fig. 11.**
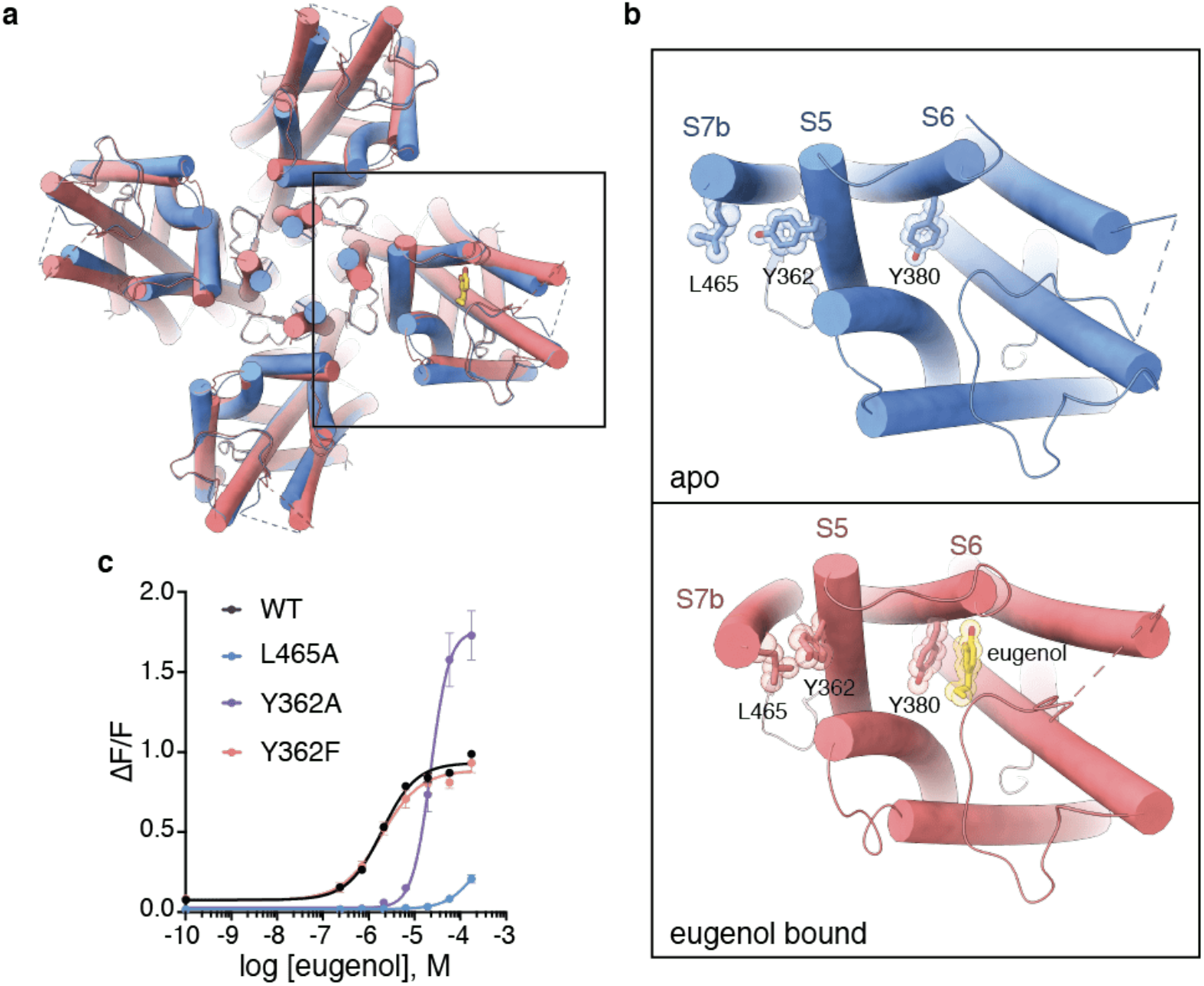
A potential route for coupling of odorant-binding to pore-opening. **a.** Top view of MhOR5 with helices represented as tubes in apo structure (blue) and eugenol-bound structure (pink). **b**. Close-up view of one subunit of the unbound (blue, top) and eugenol-bound (pink, bottom) structures of MhOR5. Residues Leu465, Tyr362 and Tyr380 and eugenol are shown as sticks with a translucent outline of the sphere representation **c**. Mutation of Leu465 in S7 and Tyr362 in S5 into alanine impairs receptor function. A conservative substitution of Tyr362 to phenylalanine restores wild-type activity, highlighting the potential role of hydrophobic packing in connecting odorant-binding with pore-opening (n = 6). Summary table of receptor data is available in Extended Data Table 6.

**Extended Data Fig. 12.**
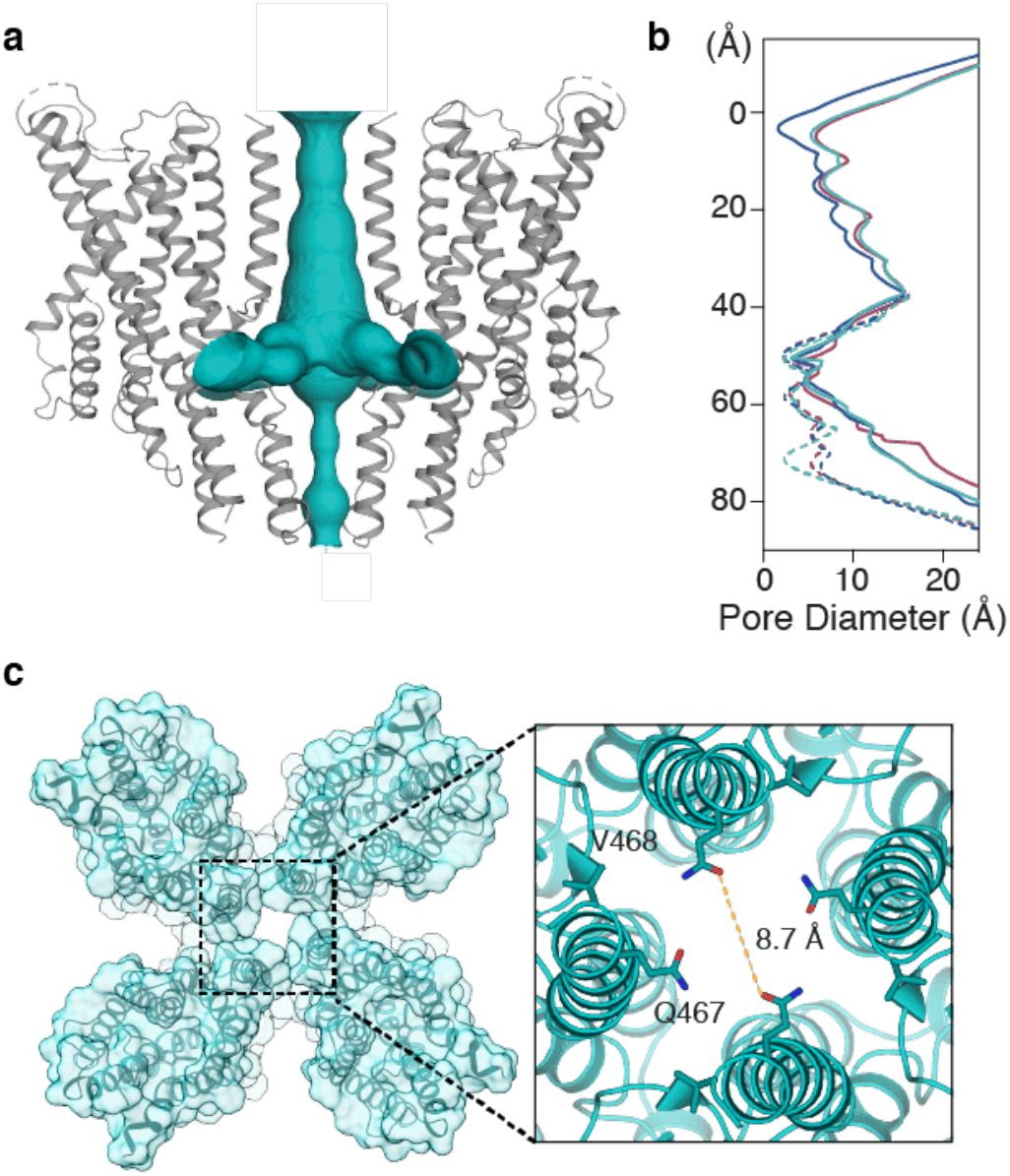
The ion permeation pathway in the DEET-bound structure of MhOR5. **a**. The lumen of the central pathway and side exits of the DEET-bound structure of MhOR5. **b**. Diameter beginning at the extracellular membrane (position ‘0’) and following the ion conduction pathway (solid line) and the central 4-fold axis through the anchor domain (dashed line), calculated with HOLE. Blue line represents unbound structure, pink line represents eugenol-bound structure and cyan line represents DEET-bound structure. **c**. Top view of the DEET-bound structure with inset (right) highlighting the positions of residue Gln467 and Val468.

**Extended Data Fig. 13.**
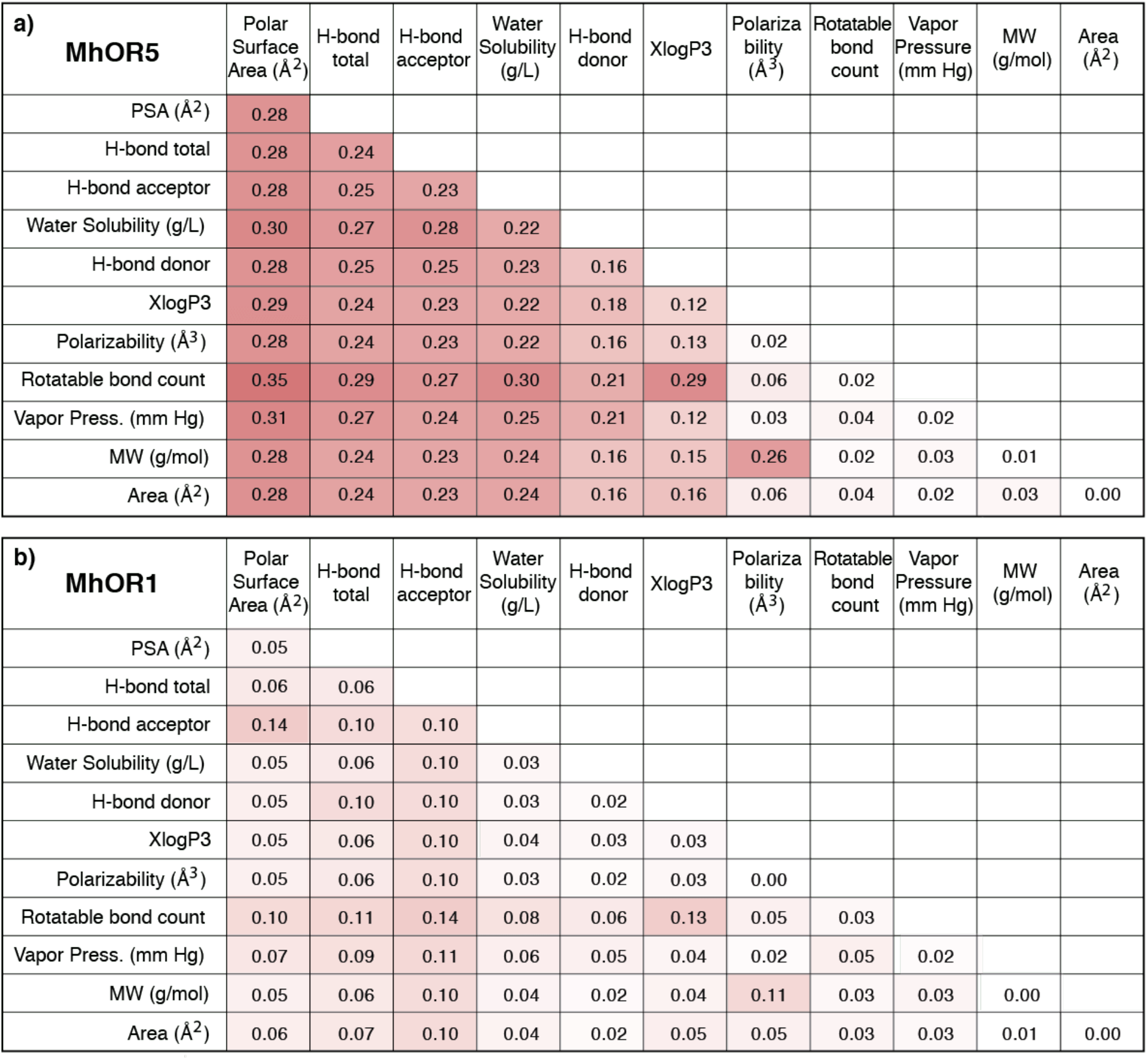
The correlation between chemical metrics and receptor activity for MhOR5 and MhOR1. **a,b**. Multiple regression analysis between pairs of chemical descriptors and receptor activity for MhOR5 (**a**) and MhOR1 (**b**). Values in each cell are the R^2^ values of the regression analysis between the descriptors in the respective column and row, combined, and the receptor’s Activity Index in response to panel of 54 ligands, colored as red-white heatmap on the same scale for all panels. Diagonal cells reflect the R^2^ values for a simple linear regression between the corresponding descriptor and Activity Index of that receptor. For both receptors, Polar Surface Area, Hydrogen Bond count, Water Solubility, Vapor Pressure, Rotatable Bond Count and Molecular Weight, individually, correlate inversely with receptor activity, and XlogP3 and Polarizability correlate positively. Note correlations are higher for MhOR5 than for MhOR1. Molecular descriptors used are compiled in Extended Data Table 10.

**Extended Data Fig. 14.**
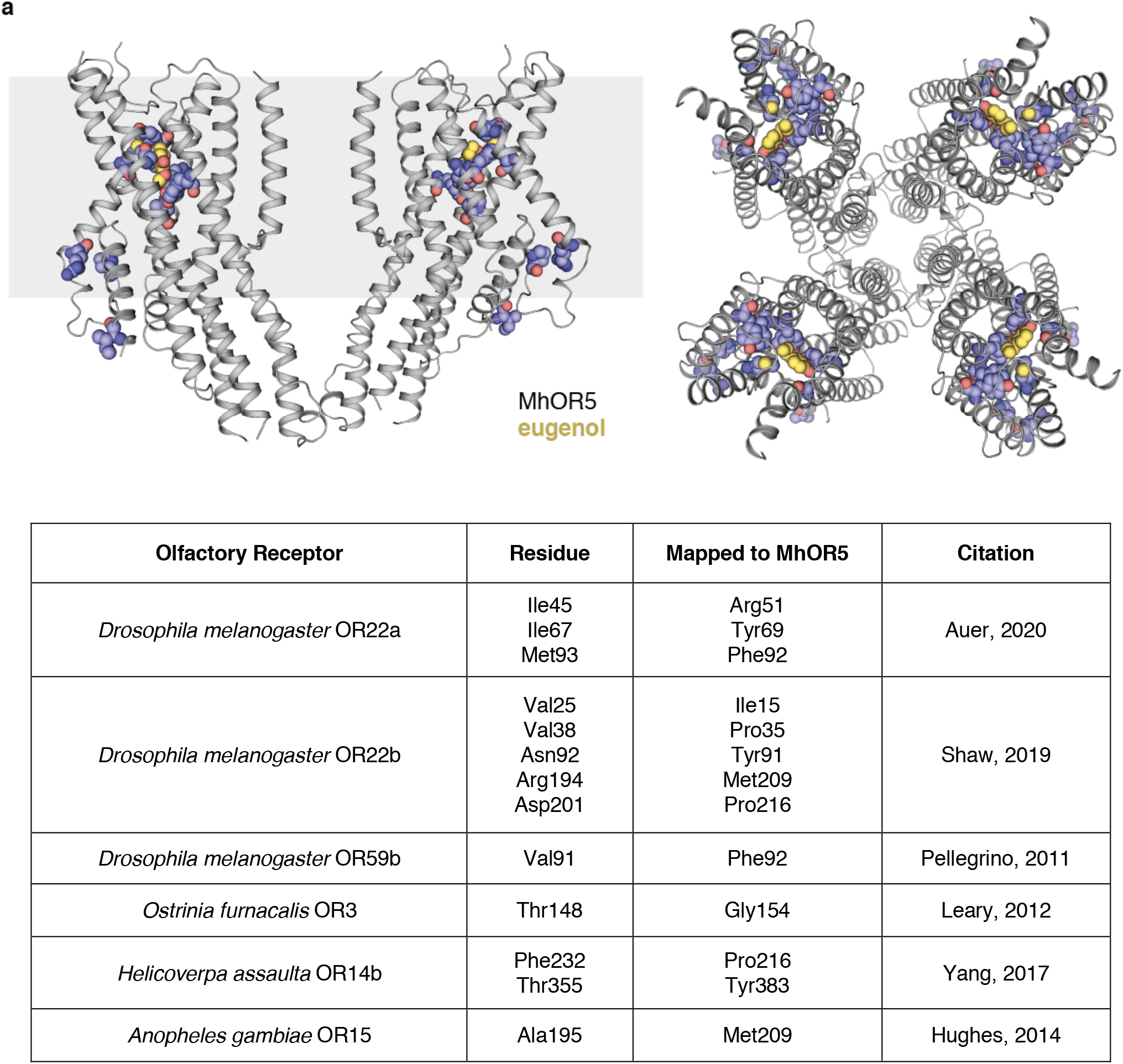
Residues implicated in odorant specificity in neopteran ORs map to the MhOR5 odorant-binding pocket. MhOR5 in ribbon representation with residues identified in from cited studies shown as spheres. Eugenol in gold, shown as spheres. Table lists amino-acid residues from the literature, their analogous position in MhOR5, and the citation of the original article. Sequence alignment was made using the PROMALS3D server (see Methods).

**Extended Data Table 1.** Cryo-EM data collection, refinement and model statistics.

|  | **Apo** | **Eugenol** | **DEET** |
| --- | --- | --- | --- |
| **Data deposition** |  |  |  |
| PDB | xxxx | xxxx | xxxx |
| EMDB |  |  |  |
| **Cryo-EM Data Collection** |  |  |  |
| Voltage (kV) | 300 | 300 | 300 |
| Magnification (x) | 29000 | 29000 | 29000 |
| Pixel Size (Å) | 1.03 | 1.03 | 1.03 |
| Electron Exposure (e-/Å<sup>2</sup>/frame) | 1.5 | 1.22 | 1.3 |
| Defocus Range (μm) | -1 to -3 | -1 to -3 | -1 to -3 |
| **Cryo-EM Data Processing** |  |  |  |
| Number of particles | 49,832 | 53,900 | 56,190 |
| Symmetry imposed | C4 | C4 | C4 |
| Map Sharpening b-factor (Å<sup>2</sup>) | -121.8 | -99 | -99.9 |
| Map Resolution (Å) | 3.3 | 2.9 | 2.9 |
| Map Resolution Range (Å) | 3.3-4.5 | 2.8-4.5 | 2.9-5 |
| FSC Threshold | 0.143 | 0.143 | 0.143 |
| **Model Refinement** |  |  |  |
| Number of MhOR5 Aminoacids | 377 | 381 | 381 |
| Number of Ligands | 0 | 1 | 1 |
| Average b-factor (Å<sup>2</sup>) | 121.19 | 39.04 | 59.08 |
| Bond Length rmsd (Å) | 0.004 | 0.005 | 0.005 |
| Bond Angles rmsd (Å) | 0.8 | 0.833 | 0.785 |
| Ramachandran Plot: |  |  |  |
| Favored (%) | 98.64 | 96.53 | 97.07 |
| Allowed (%) | 1.36 | 3.47 | 2.93 |
| Outliers (%) | 0 | 0 | 0 |
| Rotamer Outliers (%) | 0.9 | 1.47 | 1.47 |
| MolProbity Score | 1.28 | 1.5 | 1.5 |
| MolProbity Clash Score | 5.27 | 3.54 | 4.28 |

**Extended Data Table 2.** MhOR5 receptor response to a panel of odorants, tastants and synthetic ligands, assayed in the functional GCaMP assay (Fig. **1c**, **4e**, Extended Data Fig. **1d-g**, **2**). Activity Index, log(EC_50_) and max ΔF/F are three metrics used to characterized the dose-response curve for each ligand. Dose-response curves that did not saturate according to the fitted Hill equation were assigned a log(EC_50_) value of −2. The Activity Index is defined as the negative product of log(EC_50_) and max ΔF/F. All values are shown with SEM.

| Ligand | CAS # | Activity Index | log(EC<sub>50</sub>) | max ΔF/F | N |
| --- | --- | --- | --- | --- | --- |
| 1-hexanol | 111-27-3 | 4.202 ± 0.156 | -4.324 ± 0.047 | 0.972 ± 0.036 | 10 |
| 1-octanol | 111-87-5 | 4.661 ± 0.140 | -4.985 ± 0.033 | 0.935 ± 0.026 | 13 |
| (R)-1-octen-3-ol | 3391-86-4 | 4.692 ± 0.324 | -4.931 ± 0.131 | 0.953 ± 0.061 | 12 |
| 1-pentanol | 71-41-0 | 1.434 ± 0.062 | -2.000 ± 0.000 | 0.717 ± 0.031 | 11 |
| 2-acetylthiophene | 88-15-3 | 4.545 ± 0.178 | -5.245 ± 0.079 | 0.868 ± 0.034 | 14 |
| 2-ethylphenol | 90-00-6 | 5.143 ± 0.211 | -5.062 ± 0.109 | 1.015 ± 0.033 | 15 |
| 2-heptanone | 110-43-0 | 4.733 ± 0.233 | -4.956 ± 0.080 | 0.955 ± 0.045 | 11 |
| 2-undecanone | 112-12-9 | 0.437 ± 0.082 | -2.000 ± 0.000 | 0.218 ± 0.041 | 7 |
| 2,3-butanediol | 513-85-9 | 0.357 ± 0.061 | -2.000 ± 0.000 | 0.178 ± 0.030 | 3 |
| 2,4,5-trimethylthiazole | 13623-11-5 | 4.641 ± 0.173 | -4.875 ± 0.062 | 0.954 ± 0.038 | 15 |
| 3-octanol | 589-98-0 | 4.856 ± 0.144 | -5.018 ± 0.045 | 0.968 ± 0.029 | 13 |
| 4-ethylphenol | 123-07-9 | 5.177 ± 0.202 | -5.158 ± 0.070 | 1.003 ± 0.036 | 15 |
| 4-methoxyphenylacetone | 122-84-9 | 4.867 ± 0.188 | -4.887 ± 0.079 | 0.994 ± 0.030 | 13 |
| acetaldehyde diethyl acetal | 105-57-7 | 1.169 ± 0.133 | -2.000 ± 0.000 | 0.584 ± 0.067 | 7 |
| acetic acid | 64-19-7 | 0.265 ± 0.081 | -2.000 ± 0.000 | 0.133 ± 0.040 | 4 |
| acetophenone | 98-86-2 | 5.265 ± 0.162 | -5.487 ± 0.052 | 0.961 ± 0.031 | 13 |
| alpha-pinene | 7785-70-8 | 4.188 ± 0.151 | -4.314 ± 0.047 | 0.972 ± 0.036 | 11 |
| L-(+)-arabinose | 5328-37-0 | 0.202 ± 0.062 | -2.000 ± 0.000 | 0.101 ± 0.031 | 4 |
| benzaldehyde | 100-52-7 | 5.314 ± 0.291 | -4.976 ± 0.080 | 1.068 ± 0.055 | 13 |
| butyl acetate | 123-86-4 | 5.107 ± 0.273 | -5.047 ± 0.068 | 1.014 ± 0.054 | 13 |
| butyric acid | 107-92-6 | 0.500 ± 0.088 | -2.000 ± 0.000 | 0.250 ± 0.044 | 11 |
| caffeine | 58-08-2 | 0.108 ± 0.043 | -2.000 ± 0.000 | 0.054 ± 0.021 | 4 |
| citric acid | 77-92-9 | 0.175 ± 0.054 | -2.000 ± 0.000 | 0.087 ± 0.027 | 4 |
| decanal | 112-31-2 | 1.399 ± 0.063 | -2.000 ± 0.000 | 0.700 ± 0.032 | 11 |
| DEET | 134-62-3 | 3.114 ± 0.155 | -4.298 ± 0.061 | 0.724 ± 0.035 | 12 |
| denatonium benzoate | 3734-33-6 | -0.035 ± 0.012 | -3.963* ± 0.056 | -0.009 ± 0.003 | 7 |
| ethyl acetate | 141-78-6 | 0.707 ± 0.096 | -2.000 ± 0.000 | 0.354 ± 0.048 | 7 |
| ethyl butyrate | 105-54-4 | 4.161 ± 0.191 | -4.147 ± 0.114 | 1.006 ± 0.041 | 14 |
| ethyl hexanoate | 123-66-0 | 5.060 ± 0.191 | -5.129 ± 0.091 | 0.983 ± 0.027 | 17 |
| eugenol | 97-53-0 | 5.701 ± 0.026 | -5.702 ± 0.018 | 1.000 ± 0.000 | 122 |
| (±)-geosmin | 16423-19-1 | 3.677 ± 0.162 | -4.308 ± 0.071 | 0.855 ± 0.040 | 11 |
| D-(+)-glucose | 50-99-7 | 0.232 ± 0.101 | -2.000 ± 0.000 | 0.116 ± 0.051 | 4 |
| heptanoic acid | 111-14-8 | 0.562 ± 0.086 | -2.000 ± 0.000 | 0.281 ± 0.043 | 12 |
| hexanal | 66-25-1 | 4.228 ± 0.249 | -4.480 ± 0.097 | 0.943 ± 0.051 | 11 |
| indole | 120-72-9 | 5.053 ± 0.280 | -5.106 ± 0.139 | 0.982 ± 0.036 | 13 |
| isobutyl acetate | 110-19-0 | 4.552 ± 0.159 | -4.705 ± 0.076 | 0.967 ± 0.030 | 14 |
| isopropyl tiglate | 1733-25-1 | 4.357 ± 0.180 | -4.669 ± 0.061 | 0.934 ± 0.039 | 13 |
| L-(+)-lactic acid | 79-33-4 | 0.419 ± 0.094 | -2.000 ± 0.000 | 0.210 ± 0.047 | 10 |
| (R)-(+)-limonene | 5989-27-5 | 4.964 ± 0.141 | -5.539 ± 0.115 | 0.899 ± 0.025 | 13 |
| linalool | 78-70-6 | 4.111 ± 0.139 | -4.572 ± 0.053 | 0.899 ± 0.026 | 14 |
| methyl benzoate | 93-58-3 | 5.420 ± 0.334 | -5.348 ± 0.124 | 1.010 ± 0.052 | 13 |
| methyl hexanoate | 106-70-7 | 4.907 ± 0.252 | -5.075 ± 0.112 | 0.967 ± 0.043 | 14 |
| methyl laurate | 111-82-0 | 0.346 ± 0.064 | -2.000 ± 0.000 | 0.173 ± 0.032 | 11 |
| L-glutamic acid monosodium salt monohydrate (MSG) | 6106-04-3 | 0.283 ± 0.092 | -2.000 ± 0.000 | 0.141 ± 0.046 | 4 |
| n-caproic acid | 142-62-1 | 0.641 ± 0.140 | -2.000 ± 0.000 | 0.321 ± 0.070 | 7 |
| o-cresol | 95-48-7 | 4.277 ± 0.164 | -4.550 ± 0.053 | 0.942 ± 0.038 | 13 |
| octanoic acid | 124-07-2 | 0.794 ± 0.097 | -2.000 ± 0.000 | 0.397 ± 0.049 | 12 |
| prenyl acetate | 1191-16-8 | 4.197 ± 0.174 | -4.542 ± 0.071 | 0.928 ± 0.043 | 12 |
| propyl acetate | 109-60-4 | 4.103 ± 0.207 | -4.198 ± 0.046 | 0.977 ± 0.047 | 11 |
| sucrose | 57-50-1 | 0.215 ± 0.061 | -2.000 ± 0.000 | 0.108 ± 0.030 | 4 |
| sulcatone | 110-93-0 | 4.682 ± 0.228 | -4.773 ± 0.060 | 0.982 ± 0.047 | 13 |
| thiazole | 288-47-1 | 0.899 ± 0.097 | -2.000 ± 0.000 | 0.450 ± 0.049 | 10 |
| *trans*-3-hexen-1-ol | 928-97-2 | 1.719 ± 0.054 | -2.000 ± 0.000 | 0.859 ± 0.027 | 10 |
| VUAA1 | 525582-84-7 | -0.105 ± 0.053 | -4.814* ± 0.141 | -0.022 ± 0.012 | 7 |

**Extended Data Table 3.** Ordering of compounds in MhOR5 tuning curves in Extended Data Fig. **2a-c**.

| Order | a) Activity | b) $\Delta F/F$ | c) - log($EC_{50}$) |
| --- | --- | --- | --- |
| 1 | VUAA1 | VUAA1 | 2,3-butanediol |
| 2 | caffeine | caffeine | acetic acid |
| 3 | L-(+)-arabinose | L-(+)-arabinose | methyl laurate |
| 4 | D-(+)-glucose | D-(+)-glucose | butyric acid |
| 5 | L-glutamic acid monosodium salt monohydrate (MSG) | L-glutamic acid monosodium salt monohydrate (MSG) | citric acid |
| 6 | methyl laurate | methyl laurate | ethyl acetate |
| 7 | 2-undecanone | 2-undecanone | heptanoic acid |
| 8 | heptanoic acid | heptanoic acid | n-caproic acid |
| 9 | ethyl acetate | ethyl acetate | sucrose |
| 10 | thiazole | thiazole | 1-pentanol |
| 11 | 1-pentanol | 1-pentanol | *trans*-3-hexen-1-ol |
| 12 | *trans*-3-hexen-1-ol | (±)-geosmin | propyl acetate |
| 13 | (±)-geosmin | *trans*-3-hexen-1-ol | (±)-geosmin |
| 14 | linalool | linalool | 1-hexanol |
| 15 | alpha-pinene | prenyl acetate | prenyl acetate |
| 16 | 1-hexanol | isopropyl tiglate | linalool |
| 17 | o-cresol | hexanal | isobutyl acetate |
| 18 | 2-acetylthiophene | 2,4,5-trimethylthiazole | 2,4,5-trimethylthiazole |
| 19 | 2,4,5-trimethylthiazole | acetophenone | (R)-1-octen-3-ol |
| 20 | sulcatone | isobutyl acetate | benzaldehyde |
| 21 | 2-heptanone | alpha-pinene | 3-octanol |
| 22 | 4-methoxyphenylacetone | propyl acetate | 2-ethylphenol |
| 23 | (R)-(+)-limonene | indole | indole |
| 24 | ethyl hexanoate | 4-methoxyphenylacetone | 4-ethylphenol |
| 25 | 2-ethylphenol | eugenol | methyl benzoate |
| 26 | acetophenone | methyl benzoate | (R)-(+)-limonene |
| 27 | methyl benzoate | 2-ethylphenol | eugenol |
| 28 | eugenol | benzaldehyde | acetophenone |
| 29 | benzaldehyde | butyl acetate | 2-acetylthiophene |
| 30 | 4-ethylphenol | ethyl butyrate | ethyl hexanoate |
| 31 | butyl acetate | 4-ethylphenol | methyl hexanoate |
| 32 | indole | ethyl hexanoate | butyl acetate |
| 33 | methyl hexanoate | sulcatone | 1-octanol |
| 34 | 3-octanol | 1-hexanol | 2-heptanone |
| 35 | (R)-1-octen-3-ol | 3-octanol | 4-methoxyphenylacetone |
| 36 | 1-octanol | methyl hexanoate | sulcatone |
| 37 | isobutyl acetate | 2-heptanone | isopropyl tiglate |
| 38 | isopropyl tiglate | (R)-1-octen-3-ol | o-cresol |
| 39 | hexanal | o-cresol | hexanal |
| 40 | prenyl acetate | 1-octanol | alpha-pinene |
| 41 | ethyl butyrate | (R)-(+)-limonene | DEET |
| 42 | propyl acetate | 2-acetylthiophene | ethyl butyrate |
| 43 | DEET | DEET | 2-undecanone |
| 44 | decanal | decanal | acetaldehyde diethyl acetal |
| 45 | acetaldehyde diethyl acetal | acetaldehyde diethyl acetal | L-(+)-arabinose |
| 46 | octanoic acid | octanoic acid | caffeine |
| 47 | n-caproic acid | n-caproic acid | decanal |
| 48 | butyric acid | butyric acid | L-(+)-lactic acid |
| 49 | L-(+)-lactic acid | L-(+)-lactic acid | D-(+)-glucose |
| 50 | 2,3-butanediol | 2,3-butanediol | L-glutamic acid monosodium salt monohydrate (MSG) |
| 51 | acetic acid | acetic acid | octanoic acid |
| 52 | sucrose | sucrose | thiazole |
| 53 | citric acid | citric acid | denatonium benzoate |
| 54 | denatonium benzoate | denatonium benzoate | VUAA1 |

**Extended Data Table 4.**
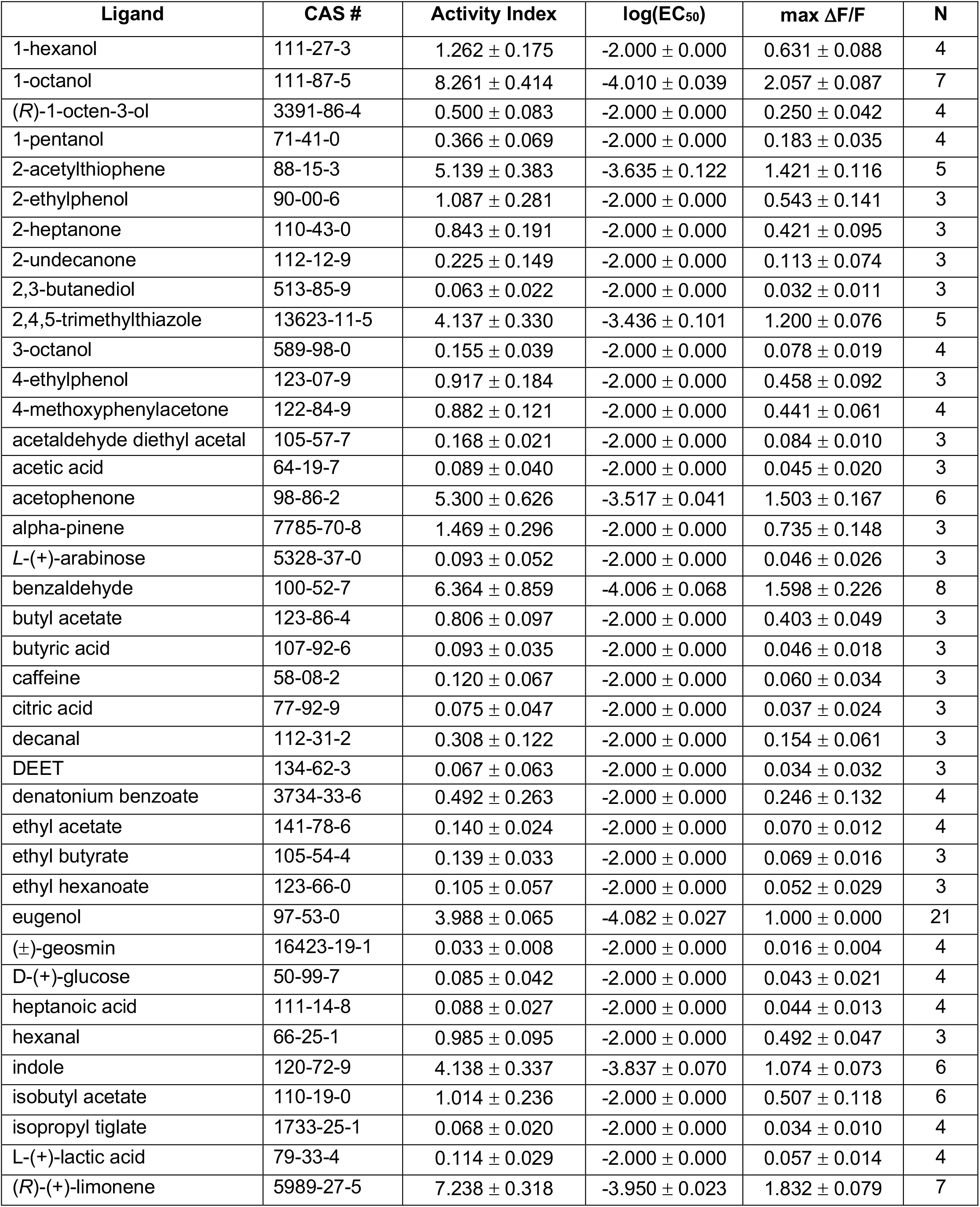
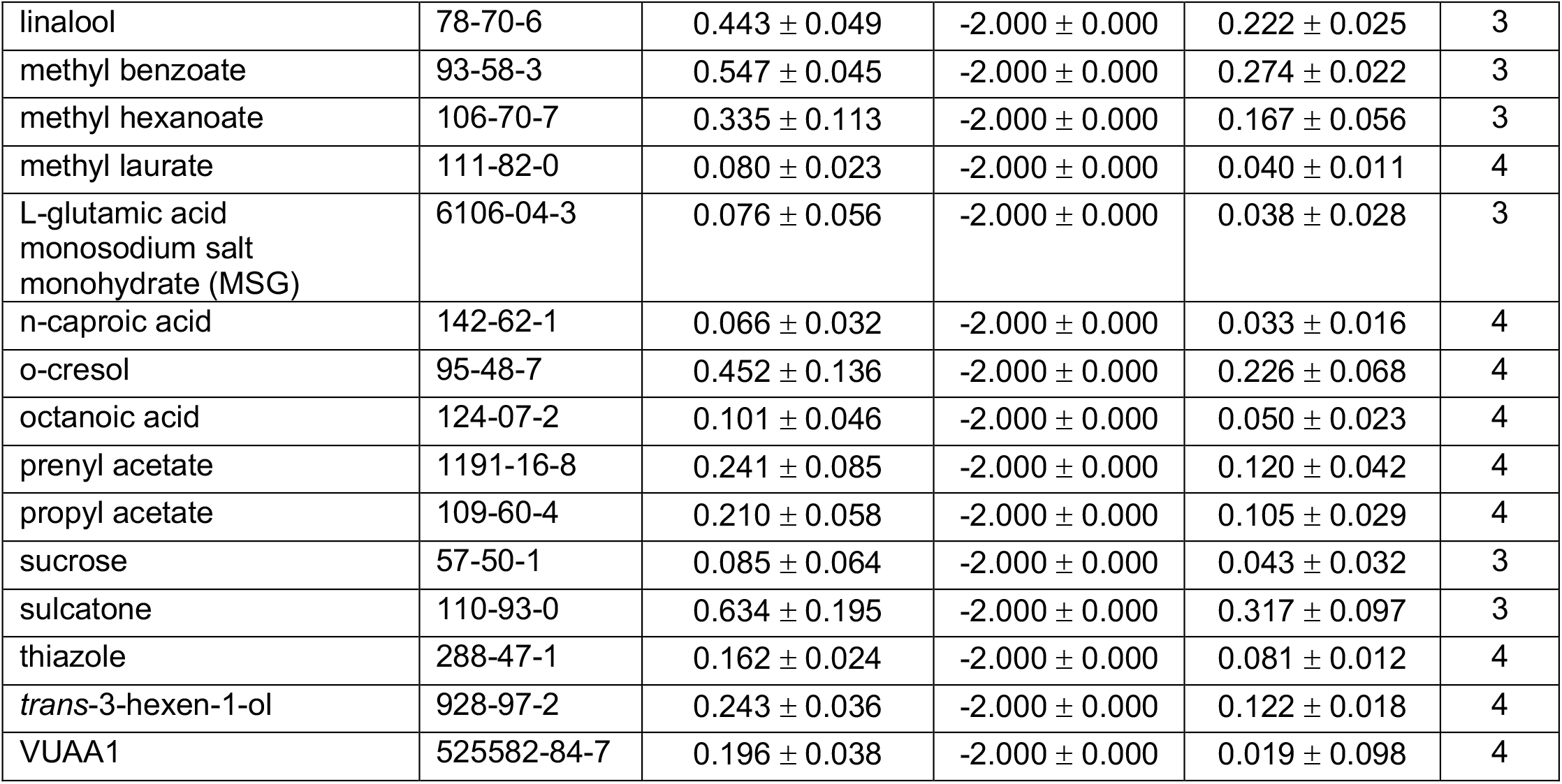
Response of MhOR1 to a panel of odorants, tastants and synthetic ligands in the functional GCaMP assay (Extended Data Fig. 1**d-g**). The Activity Index is defined as the negative product of log(EC_50_) and max ΔF/F. Dose-response curves that did not saturate according to the fitted Hill equation were assigned a log(EC_50_) value of −2. All values are shown with SEM.

| Ligand | CAS # | Activity Index | $\log(\text{EC}_{50})$ | $\max \Delta F/F$ | N |
| --- | --- | --- | --- | --- | --- |
| 1-hexanol | 111-27-3 | $1.262 \pm 0.175$ | $-2.000 \pm 0.000$ | $0.631 \pm 0.088$ | 4 |
| 1-octanol | 111-87-5 | $8.261 \pm 0.414$ | $-4.010 \pm 0.039$ | $2.057 \pm 0.087$ | 7 |
| (R)-1-octen-3-ol | 3391-86-4 | $0.500 \pm 0.083$ | $-2.000 \pm 0.000$ | $0.250 \pm 0.042$ | 4 |
| 1-pentanol | 71-41-0 | $0.366 \pm 0.069$ | $-2.000 \pm 0.000$ | $0.183 \pm 0.035$ | 4 |
| 2-acetylthiophene | 88-15-3 | $5.139 \pm 0.383$ | $-3.635 \pm 0.122$ | $1.421 \pm 0.116$ | 5 |
| 2-ethylphenol | 90-00-6 | $1.087 \pm 0.281$ | $-2.000 \pm 0.000$ | $0.543 \pm 0.141$ | 3 |
| 2-heptanone | 110-43-0 | $0.843 \pm 0.191$ | $-2.000 \pm 0.000$ | $0.421 \pm 0.095$ | 3 |
| 2-undecanone | 112-12-9 | $0.225 \pm 0.149$ | $-2.000 \pm 0.000$ | $0.113 \pm 0.074$ | 3 |
| 2,3-butanediol | 513-85-9 | $0.063 \pm 0.022$ | $-2.000 \pm 0.000$ | $0.032 \pm 0.011$ | 3 |
| 2,4,5-trimethylthiazole | 13623-11-5 | $4.137 \pm 0.330$ | $-3.436 \pm 0.101$ | $1.200 \pm 0.076$ | 5 |
| 3-octanol | 589-98-0 | $0.155 \pm 0.039$ | $-2.000 \pm 0.000$ | $0.078 \pm 0.019$ | 4 |
| 4-ethylphenol | 123-07-9 | $0.917 \pm 0.184$ | $-2.000 \pm 0.000$ | $0.458 \pm 0.092$ | 3 |
| 4-methoxyphenylacetone | 122-84-9 | $0.882 \pm 0.121$ | $-2.000 \pm 0.000$ | $0.441 \pm 0.061$ | 4 |
| acetaldehyde diethyl acetal | 105-57-7 | $0.168 \pm 0.021$ | $-2.000 \pm 0.000$ | $0.084 \pm 0.010$ | 3 |
| acetic acid | 64-19-7 | $0.089 \pm 0.040$ | $-2.000 \pm 0.000$ | $0.045 \pm 0.020$ | 3 |
| acetophenone | 98-86-2 | $5.300 \pm 0.626$ | $-3.517 \pm 0.041$ | $1.503 \pm 0.167$ | 6 |
| alpha-pinene | 7785-70-8 | $1.469 \pm 0.296$ | $-2.000 \pm 0.000$ | $0.735 \pm 0.148$ | 3 |
| L-(+)-arabinose | 5328-37-0 | $0.093 \pm 0.052$ | $-2.000 \pm 0.000$ | $0.046 \pm 0.026$ | 3 |
| benzaldehyde | 100-52-7 | $6.364 \pm 0.859$ | $-4.006 \pm 0.068$ | $1.598 \pm 0.226$ | 8 |
| butyl acetate | 123-86-4 | $0.806 \pm 0.097$ | $-2.000 \pm 0.000$ | $0.403 \pm 0.049$ | 3 |
| butyric acid | 107-92-6 | $0.093 \pm 0.035$ | $-2.000 \pm 0.000$ | $0.046 \pm 0.018$ | 3 |
| caffeine | 58-08-2 | $0.120 \pm 0.067$ | $-2.000 \pm 0.000$ | $0.060 \pm 0.034$ | 3 |
| citric acid | 77-92-9 | $0.075 \pm 0.047$ | $-2.000 \pm 0.000$ | $0.037 \pm 0.024$ | 3 |
| decanal | 112-31-2 | $0.308 \pm 0.122$ | $-2.000 \pm 0.000$ | $0.154 \pm 0.061$ | 3 |
| DEET | 134-62-3 | $0.067 \pm 0.063$ | $-2.000 \pm 0.000$ | $0.034 \pm 0.032$ | 3 |
| denatonium benzoate | 3734-33-6 | $0.492 \pm 0.263$ | $-2.000 \pm 0.000$ | $0.246 \pm 0.132$ | 4 |
| ethyl acetate | 141-78-6 | $0.140 \pm 0.024$ | $-2.000 \pm 0.000$ | $0.070 \pm 0.012$ | 4 |
| ethyl butyrate | 105-54-4 | $0.139 \pm 0.033$ | $-2.000 \pm 0.000$ | $0.069 \pm 0.016$ | 3 |
| ethyl hexanoate | 123-66-0 | $0.105 \pm 0.057$ | $-2.000 \pm 0.000$ | $0.052 \pm 0.029$ | 3 |
| eugenol | 97-53-0 | $3.988 \pm 0.065$ | $-4.082 \pm 0.027$ | $1.000 \pm 0.000$ | 21 |
| (±)-geosmin | 16423-19-1 | $0.033 \pm 0.008$ | $-2.000 \pm 0.000$ | $0.016 \pm 0.004$ | 4 |
| D-(+)-glucose | 50-99-7 | $0.085 \pm 0.042$ | $-2.000 \pm 0.000$ | $0.043 \pm 0.021$ | 4 |
| heptanoic acid | 111-14-8 | $0.088 \pm 0.027$ | $-2.000 \pm 0.000$ | $0.044 \pm 0.013$ | 4 |
| hexanal | 66-25-1 | $0.985 \pm 0.095$ | $-2.000 \pm 0.000$ | $0.492 \pm 0.047$ | 3 |
| indole | 120-72-9 | $4.138 \pm 0.337$ | $-3.837 \pm 0.070$ | $1.074 \pm 0.073$ | 6 |
| isobutyl acetate | 110-19-0 | $1.014 \pm 0.236$ | $-2.000 \pm 0.000$ | $0.507 \pm 0.118$ | 6 |
| isopropyl tiglate | 1733-25-1 | $0.068 \pm 0.020$ | $-2.000 \pm 0.000$ | $0.034 \pm 0.010$ | 4 |
| L-(+)-lactic acid | 79-33-4 | $0.114 \pm 0.029$ | $-2.000 \pm 0.000$ | $0.057 \pm 0.014$ | 4 |
| (R)-(+)-limonene | 5989-27-5 | $7.238 \pm 0.318$ | $-3.950 \pm 0.023$ | $1.832 \pm 0.079$ | 7 |
| linalool | 78-70-6 | 0.443 ± 0.049 | -2.000 ± 0.000 | 0.222 ± 0.025 | 3 |
| methyl benzoate | 93-58-3 | 0.547 ± 0.045 | -2.000 ± 0.000 | 0.274 ± 0.022 | 3 |
| methyl hexanoate | 106-70-7 | 0.335 ± 0.113 | -2.000 ± 0.000 | 0.167 ± 0.056 | 3 |
| methyl laurate | 111-82-0 | 0.080 ± 0.023 | -2.000 ± 0.000 | 0.040 ± 0.011 | 4 |
| L-glutamic acid monosodium salt monohydrate (MSG) | 6106-04-3 | 0.076 ± 0.056 | -2.000 ± 0.000 | 0.038 ± 0.028 | 3 |
| n-caproic acid | 142-62-1 | 0.066 ± 0.032 | -2.000 ± 0.000 | 0.033 ± 0.016 | 4 |
| o-cresol | 95-48-7 | 0.452 ± 0.136 | -2.000 ± 0.000 | 0.226 ± 0.068 | 4 |
| octanoic acid | 124-07-2 | 0.101 ± 0.046 | -2.000 ± 0.000 | 0.050 ± 0.023 | 4 |
| prenyl acetate | 1191-16-8 | 0.241 ± 0.085 | -2.000 ± 0.000 | 0.120 ± 0.042 | 4 |
| propyl acetate | 109-60-4 | 0.210 ± 0.058 | -2.000 ± 0.000 | 0.105 ± 0.029 | 4 |
| sucrose | 57-50-1 | 0.085 ± 0.064 | -2.000 ± 0.000 | 0.043 ± 0.032 | 3 |
| sulcatone | 110-93-0 | 0.634 ± 0.195 | -2.000 ± 0.000 | 0.317 ± 0.097 | 3 |
| thiazole | 288-47-1 | 0.162 ± 0.024 | -2.000 ± 0.000 | 0.081 ± 0.012 | 4 |
| *trans*-3-hexen-1-ol | 928-97-2 | 0.243 ± 0.036 | -2.000 ± 0.000 | 0.122 ± 0.018 | 4 |
| VUAA1 | 525582-84-7 | 0.196 ± 0.038 | -2.000 ± 0.000 | 0.019 ± 0.098 | 4 |

**Extended Data Table 5.** Lifetime sparseness values for *M. hrabei* OR5 and OR1 and *D. melanogaster* ORs from published data (Extended Data Fig. **1e**). Those values closest to 0 correspond to broadly tuned receptors that respond similarly to many ligands in a set, while values closest to 1 are narrowly tuned to a single or to a small subset of ligands. All inhibitory responses to odorants set to 0 before calculation.

| Odorant Receptor | # odorants | Lifetime Sparseness Value |
| --- | --- | --- |
| *M. hrabei* OR5 | 54 | 0.346 |
| *M. hrabei* OR1 | 54 | 0.764 |
| *D. melanogaster* OR1a | 30 | 0.458 |
| *D. melanogaster* OR2a | 124 | 0.607 |
| *D. melanogaster* OR7a | 222 | 0.657 |
| *D. melanogaster* OR9a | 144 | 0.546 |
| *D. melanogaster* OR10a | 235 | 0.771 |
| *D. melanogaster* OR13a | 167 | 0.606 |
| *D. melanogaster* OR19a | 497 | 0.817 |
| *D. melanogaster* OR22a | 225 | 0.585 |
| *D. melanogaster* OR22b | 11 | 0.526 |
| *D. melanogaster* OR23a | 115 | 0.757 |
| *D. melanogaster* OR30a | 30 | 0.811 |
| *D. melanogaster* OR33a | 30 | 0.280 |
| *D. melanogaster* OR33b | 122 | 0.761 |
| *D. melanogaster* OR33c | 12 | 0.472 |
| *D. melanogaster* OR35a | 123 | 0.673 |
| *D. melanogaster* OR42a | 72 | 0.702 |
| *D. melanogaster* OR42b | 177 | 0.795 |
| *D. melanogaster* OR43a | 115 | 0.793 |
| *D. melanogaster* OR43b | 144 | 0.670 |
| *D. melanogaster* OR45a | 31 | 0.480 |
| *D. melanogaster* OR45b | 30 | 0.812 |
| *D. melanogaster* OR46a | 12 | 0.983 |
| *D. melanogaster* OR47a | 135 | 0.716 |
| *D. melanogaster* OR47b | 178 | 0.872 |
| *D. melanogaster* OR49a | 30 | 0.477 |
| *D. melanogaster* OR49b | 164 | 0.880 |
| *D. melanogaster* OR59a | 29 | 0.662 |
| *D. melanogaster* OR59b | 173 | 0.810 |
| *D. melanogaster* OR59c | 53 | 0.697 |
| *D. melanogaster* OR65a | 116 | 0.754 |
| *D. melanogaster* OR67a | 127 | 0.583 |
| *D. melanogaster* OR67b | 121 | 0.522 |
| *D. melanogaster* OR67c | 161 | 0.716 |
| *D. melanogaster* OR69a | 107 | 0.149 |
| *D. melanogaster* OR71a | 149 | 0.790 |
| *D. melanogaster* OR74a | 30 | 0.619 |
| *D. melanogaster* OR82a | 180 | 0.808 |
| *D. melanogaster* OR83c | 125 | 0.667 |
| *D. melanogaster* OR85a | 114 | 0.850 |
| *D. melanogaster* OR85b | 161 | 0.620 |
| *D. melanogaster* OR85c | 30 | 0.622 |
| *D. melanogaster* OR85d | 51 | 0.809 |
| *D. melanogaster* OR85e | 12 | 0.531 |
| *D. melanogaster* OR85f | 114 | 0.473 |
| *D. melanogaster* OR88a | 115 | 0.878 |
| *D. melanogaster* OR92a | 174 | 0.853 |
| *D. melanogaster* OR94a | 30 | 0.863 |
| *D. melanogaster* OR94b | 31 | 0.602 |
| *D. melanogaster* OR98a | 161 | 0.673 |

**Extended Data Table 6.** Response of wild-type and mutant MhOR5 receptors to eugenol, assayed in the functional GCaMP assay (Fig. **2e**, **3d**, **4d**, Extended Data Fig. **10**, **11c**). Baseline fluorescence is normalized to wild-type MhOR5 baseline fluorescence on the same plate. Dose-response curves that did not saturate according to the fitted Hill equation were assigned a log(EC_50_) value of −2. The Activity Index is defined as the negative product of log(EC_50_) and max ΔF/F. All values are shown with SEM.

| Mutation | Normalized Baseline | Activity Index | log(EC<sub>50</sub>) | max ΔF/F | N |
| --- | --- | --- | --- | --- | --- |
| wild-type MhOR5 | 1.000 ± 0.000 | 5.701 ± 0.026 | -5.702 ± 0.018 | 1.000 ± 0.000 | 122 |
| T87A | 0.764 ± 0.036 | 6.635 ± 0.385 | -5.590 ± 0.058 | 1.186 ± 0.064 | 7 |
| V88A | 0.712 ± 0.037 | 6.817 ± 0.503 | -4.907 ± 0.048 | 1.391 ± 0.108 | 6 |
| Y91A | 0.516 ± 0.034 | 9.811 ± 0.588 | -4.589 ± 0.061 | 2.133 ± 0.107 | 6 |
| F92A | 1.894 ± 0.083 | 0.545 ± 0.035 | -2.000 ± 0.000 | 0.272 ± 0.018 | 7 |
| S151A | 2.976 ± 0.114 | 0.098 ± 0.010 | -2.000 ± 0.000 | 0.049 ± 0.005 | 6 |
| G154A | 1.356 ± 0.093 | 0.583 ± 0.053 | -2.000 ± 0.000 | 0.291 ± 0.026 | 6 |
| W155A | 1.279 ± 0.028 | 2.636 ± 0.131 | -5.156 ± 0.104 | 0.515 ± 0.035 | 6 |
| W158A | 0.502 ± 0.056 | 0.167 ± 0.029 | -2.000 ± 0.000 | 0.084 ± 0.014 | 6 |
| M209A | 3.044 ± 0.046 | 0.218 ± 0.047 | -2.000 ± 0.000 | 0.109 ± 0.024 | 7 |
| M209V | 0.698 ± 0.025 | 5.075 ± 0.292 | -4.707 ± 0.046 | 1.078 ± 0.062 | 6 |
| M209L | 0.663 ± 0.054 | 9.586 ± 0.873 | -5.000 ± 0.026 | 1.939 ± 0.180 | 6 |
| I213A | 2.424 ± 0.097 | 0.375 ± 0.043 | -2.000 ± 0.000 | 0.187 ± 0.022 | 6 |
| I213M | 0.952 ± 0.016 | 6.362 ± 0.865 | -5.046 ± 0.042 | 1.258 ± 0.169 | 7 |
| Y362A | 0.521 ± 0.041 | 8.120 ± 0.748 | -4.655 ± 0.031 | 1.741 ± 0.154 | 6 |
| Y362F | 0.965 ± 0.032 | 5.255 ± 0.264 | -5.606 ± 0.187 | 0.944 ± 0.061 | 6 |
| L364A | 0.528 ± 0.027 | 7.668 ± 0.303 | -4.736 ± 0.037 | 1.621 ± 0.070 | 6 |
| G376A | 1.485 ± 0.035 | 2.856 ± 0.074 | -5.573 ± 0.041 | 0.513 ± 0.015 | 7 |
| L379A | 0.831 ± 0.035 | 5.192 ± 0.305 | -5.590 ± 0.071 | 0.932 ± 0.063 | 6 |
| Y380A | 2.915 ± 0.298 | 0.115 ± 0.005 | -2.000 ± 0.000 | 0.057 ± 0.003 | 6 |
| Y383A | 0.516 ± 0.029 | 0.282 ± 0.035 | -2.000 ± 0.000 | 0.141 ± 0.017 | 6 |
| L465A | 0.490 ± 0.028 | 0.441 ± 0.051 | -2.000 ± 0.000 | 0.221 ± 0.025 | 6 |
| Q467A | 0.554 ± 0.034 | 0.083 ± 0.011 | -2.000 ± 0.000 | 0.041 ± 0.005 | 7 |
| Q467E | 3.649 ± 0.247 | 0.108 ± 0.018 | -2.000 ± 0.000 | 0.054 ± 0.009 | 6 |
| Q467R | 0.594 ± 0.027 | 0.856 ± 0.185 | -2.000 ± 0.000 | 0.428 ± 0.092 | 7 |
| Q467N | 0.696 ± 0.038 | 7.810 ± 0.643 | -5.269 ± 0.137 | 1.472 ± 0.087 | 6 |
| V468A | 1.201 ± 0.037 | 4.475 ± 0.199 | -5.779 ± 0.131 | 0.778 ± 0.047 | 6 |
| V468N | 0.753 ± 0.051 | 6.725 ± 0.334 | -5.283 ± 0.079 | 1.273 ± 0.062 | 6 |

**Extended Data Table 7.** Response of wild-type and mutant *A. bakeri* Orco and Orco / *A. gambiae* OR28 tested with their cognate ligands in the functional GCaMP assay (Fig. **2f**). The Activity Index is defined as the negative product of log(EC_50_) and max ΔF/F. Dose-response curves that did not saturate according to the fitted Hill equation were assigned a log(EC_50_) value of −2. Max ΔF/F is normalized to respective wild-type heteromer. All values are shown with SEM.

| Construct | Ligand | Activity Index | $\log(\text{EC}_{50})$ | $\max \Delta F/F$ | N |
| --- | --- | --- | --- | --- | --- |
| *A. bakeri* Orco | VUAA1 | $4.258 \pm 0.030$ | $-4.258 \pm 0.030$ | $1.000 \pm 0.000$ | 8 |
| *A. bakeri* Orco Q472A | VUAA1 | $0.239 \pm 0.068$ | $-2.000 \pm 0.000$ | $0.120 \pm 0.034$ | 5 |
| *A. bakeri* Orco /<br>*A. gambiae* OR28 | acetophenone | $4.846 \pm 0.081$ | $-4.846 \pm 0.081$ | $1.000 \pm 0.000$ | 7 |
| *A. bakeri* Orco Q472A /<br>*A. gambiae* OR28 | acetophenone | $4.290 \pm 0.652$ | $-3.989 \pm 0.053$ | $1.068 \pm 0.152$ | 7 |

**Extended Data Table 8.** Response of wild-type and mutant MhOR5 receptors to DEET assayed in the functional GCaMP assay (Fig. **4d**, Extended Data Fig. **10b-d**). The Activity Index is defined as the negative product of log(EC_50_) and max ΔF/F. Max ΔF/F is normalized to wild-type MhOR5 with DEET. Dose-response curves that did not saturate according to the fitted Hill equation were assigned a log(EC_50_) value of −2. Baseline fluorescence normalized to wild-type MhOR5 baseline fluorescence on the same plate. All values are shown with SEM.

| Mutation | Normalized Baseline | Activity Index | $\log(\text{EC}_{50})$ | $\max \Delta F/F$ | N |
| --- | --- | --- | --- | --- | --- |
| wild-type MhOR5 | $1.000 \pm 0.000$ | $3.114 \pm 0.155$ | $-4.298 \pm 0.061$ | $0.724 \pm 0.035$ | 12 |
| M209A | $3.044 \pm 0.046$ | $0.895 \pm 0.122$ | $-5.243 \pm 0.100$ | $0.173 \pm 0.026$ | 9 |
| M209V | $0.698 \pm 0.025$ | $6.654 \pm 0.320$ | $-5.122 \pm 0.080$ | $1.301 \pm 0.065$ | 6 |
| M209L | $0.663 \pm 0.054$ | $6.057 \pm 1.031$ | $-4.158 \pm 0.030$ | $1.452 \pm 0.242$ | 8 |
| I213A | $2.424 \pm 0.097$ | $0.323 \pm 0.056$ | $-2.000 \pm 0.000$ | $0.162 \pm 0.028$ | 7 |
| I213M | $0.952 \pm 0.016$ | $0.042 \pm 0.007$ | $-2.000 \pm 0.000$ | $0.021 \pm 0.004$ | 6 |

**Extended Data Table 9.** **a, b**. Response of MhOR5 M209V (**a**) and MhOR5 I213M (**b**) receptors to a panel of 40 odorants in the functional GCaMP assay (Fig. **4e**, Extended Data Fig. **10b,c**). The Activity Index is defined as the negative product of log(EC_50_) and max ΔF/F. Dose-response curves that did not saturate according to the fitted Hill equation were assigned a log(EC_50_) value of −2. All values are shown with SEM. Max ΔF/F is normalized to MhOR5 with eugenol.

**a)**
| Odorant | Activity Index | $\log(\text{EC}_{50})$ | $\max \Delta F/F$ | N |
| --- | --- | --- | --- | --- |
| 1-hexanol | $1.325 \pm 0.145$ | $-2.000 \pm 0.000$ | $0.662 \pm 0.073$ | 5 |
| 1-octanol | $3.215 \pm 0.266$ | $-4.078 \pm 0.037$ | $0.789 \pm 0.066$ | 6 |
| (R)-1-octen-3-ol | $5.066 \pm 0.254$ | $-3.963 \pm 0.059$ | $1.275 \pm 0.048$ | 6 |
| 1-pentanol | $0.114 \pm 0.018$ | $-2.000 \pm 0.000$ | $0.057 \pm 0.009$ | 5 |
| 2-acetylthiophene | $2.622 \pm 0.276$ | $-3.777 \pm 0.048$ | $0.696 \pm 0.075$ | 5 |
| 2-ethylphenol | $5.261 \pm 0.327$ | $-4.376 \pm 0.023$ | $1.202 \pm 0.072$ | 6 |
| 2-heptanone | $5.141 \pm 0.308$ | $-4.073 \pm 0.064$ | $1.261 \pm 0.068$ | 6 |
| 2,4,5-trimethylthiazole | $1.733 \pm 0.141$ | $-2.000 \pm 0.000$ | $0.867 \pm 0.070$ | 6 |
| 3-octanol | $4.333 \pm 0.225$ | $-3.890 \pm 0.041$ | $1.113 \pm 0.054$ | 6 |
| 4-ethylphenol | $4.935 \pm 0.439$ | $-4.259 \pm 0.109$ | $1.158 \pm 0.101$ | 6 |
| 4-methoxyphenylacetone | $6.422 \pm 0.290$ | $-4.527 \pm 0.041$ | $1.419 \pm 0.064$ | 6 |
| acetophenone | $4.178 \pm 0.359$ | $-4.246 \pm 0.040$ | $0.983 \pm 0.081$ | 6 |
| alpha-pinene | $3.272 \pm 0.454$ | $-3.779 \pm 0.066$ | $0.864 \pm 0.115$ | 5 |
| benzaldehyde | $4.273 \pm 0.327$ | $-3.679 \pm 0.019$ | $1.161 \pm 0.088$ | 6 |
| butyl acetate | $5.069 \pm 0.439$ | $-3.903 \pm 0.054$ | $1.295 \pm 0.104$ | 6 |
| butyric acid | $0.040 \pm 0.017$ | $-2.000 \pm 0.000$ | $0.020 \pm 0.008$ | 5 |
| decanal | $0.185 \pm 0.038$ | $-2.000 \pm 0.000$ | $0.093 \pm 0.019$ | 5 |
| DEET | $6.654 \pm 0.320$ | $-5.122 \pm 0.080$ | $1.301 \pm 0.065$ | 6 |
| ethyl butyrate | $1.410 \pm 0.169$ | $-2.000 \pm 0.000$ | $0.705 \pm 0.084$ | 6 |
| ethyl hexanoate | $1.184 \pm 0.206$ | $-2.000 \pm 0.000$ | $0.592 \pm 0.103$ | 6 |
| eugenol | $5.075 \pm 0.292$ | $-4.707 \pm 0.046$ | $1.078 \pm 0.062$ | 6 |
| (±)-geosmin | $0.199 \pm 0.034$ | $-2.000 \pm 0.000$ | $0.099 \pm 0.017$ | 6 |
| heptanoic acid | $0.054 \pm 0.008$ | $-2.000 \pm 0.000$ | $0.027 \pm 0.004$ | 5 |
| hexanal | $1.536 \pm 0.187$ | $-2.000 \pm 0.000$ | $0.768 \pm 0.094$ | 5 |
| indole | $6.108 \pm 0.370$ | $-4.282 \pm 0.043$ | $1.425 \pm 0.080$ | 6 |
| isobutyl acetate | $4.824 \pm 0.387$ | $-3.765 \pm 0.033$ | $1.282 \pm 0.105$ | 6 |
| isopropyl tiglate | $4.174 \pm 0.235$ | $-3.983 \pm 0.044$ | $1.050 \pm 0.067$ | 5 |
| L-(+)-lactic acid | $0.037 \pm 0.009$ | $-2.000 \pm 0.000$ | $0.019 \pm 0.004$ | 5 |
| (R)-(+)-limonene | $5.945 \pm 0.234$ | $-5.101 \pm 0.037$ | $1.166 \pm 0.045$ | 6 |
| linalool | $3.670 \pm 0.106$ | $-3.991 \pm 0.042$ | $0.920 \pm 0.028$ | 6 |
| methyl benzoate | $3.764 \pm 0.351$ | $-4.190 \pm 0.034$ | $0.899 \pm 0.086$ | 5 |
| methyl hexanoate | $5.049 \pm 0.265$ | $-4.095 \pm 0.053$ | $1.231 \pm 0.054$ | 6 |
| methyl laurate | $0.051 \pm 0.014$ | $-2.000 \pm 0.000$ | $0.026 \pm 0.007$ | 5 |
| o-cresol | $2.036 \pm 0.116$ | $-2.000 \pm 0.000$ | $1.018 \pm 0.058$ | 6 |
| octanoic acid | $0.070 \pm 0.016$ | $-2.000 \pm 0.000$ | $-0.035 \pm 0.008$ | 5 |
| prenyl acetate | 4.806 ± 0.248 | -3.990 ± 0.024 | 1.205 ± 0.065 | 6 |
| propyl acetate | 0.864 ± 0.196 | -2.000 ± 0.000 | 0.432 ± 0.098 | 5 |
| sulcatone | 5.429 ± 0.242 | -4.226 ± 0.072 | 1.287 ± 0.063 | 6 |
| thiazole | 0.087 ± 0.014 | -2.000 ± 0.000 | 0.043 ± 0.007 | 5 |
| *trans*-3-hexen-1-ol | 0.241 ± 0.031 | -2.000 ± 0.000 | 0.120 ± 0.016 | 5 |

**b)**
| Odorant | Activity Index | log(EC<sub>50</sub>) | max ΔF/F | N |
| --- | --- | --- | --- | --- |
| 1-hexanol | 0.708 ± 0.083 | -2.000 ± 0.000 | 0.354 ± 0.041 | 5 |
| 1-octanol | 1.937 ± 0.183 | -3.870 ± 0.059 | 0.505 ± 0.055 | 6 |
| (*R*)-1-octen-3-ol | 1.588 ± 0.189 | -2.000 ± 0.000 | 0.631 ± 0.033 | 6 |
| 1-pentanol | 0.183 ± 0.023 | -2.000 ± 0.000 | 0.092 ± 0.011 | 5 |
| 2-acetylthiophene | 2.370 ± 0.181 | -4.277 ± 0.172 | 0.559 ± 0.050 | 6 |
| 2-ethylphenol | 3.661 ± 0.340 | -4.369 ± 0.102 | 0.842 ± 0.081 | 6 |
| 2-heptanone | 1.650 ± 0.094 | -2.000 ± 0.000 | 0.825 ± 0.047 | 6 |
| 2,4,5-trimethylthiazole | 2.808 ± 0.355 | -3.772 ± 0.041 | 0.754 ± 0.107 | 6 |
| 3-octanol | 1.439 ± 0.175 | -2.000 ± 0.000 | 0.632 ± 0.045 | 6 |
| 4-ethylphenol | 3.999 ± 0.252 | -4.565 ± 0.034 | 0.877 ± 0.059 | 6 |
| 4-methoxyphenylacetone | 4.724 ± 0.571 | -4.567 ± 0.101 | 1.038 ± 0.129 | 6 |
| acetophenone | 3.188 ± 0.405 | -4.349 ± 0.060 | 0.732 ± 0.090 | 6 |
| alpha-pinene | 4.274 ± 0.499 | -4.598 ± 0.096 | 0.938 ± 0.122 | 5 |
| benzaldehyde | 1.891 ± 0.188 | -2.000 ± 0.000 | 0.945 ± 0.094 | 6 |
| butyl acetate | 1.347 ± 0.583 | -3.926 ± 0.083 | 0.946 ± 0.136 | 6 |
| butyric acid | 0.065 ± 0.016 | -2.000 ± 0.000 | 0.032 ± 0.008 | 5 |
| decanal | 0.207 ± 0.031 | -2.000 ± 0.000 | 0.104 ± 0.015 | 5 |
| DEET | 0.058 ± 0.019 | -2.000 ± 0.000 | 0.021 ± 0.004 | 6 |
| ethyl butyrate | 1.107 ± 0.216 | -2.000 ± 0.000 | 0.554 ± 0.108 | 6 |
| ethyl hexanoate | 0.631 ± 0.067 | -2.000 ± 0.000 | 0.315 ± 0.034 | 6 |
| eugenol | 6.362 ± 0.865 | -5.046 ± 0.042 | 1.258 ± 0.169 | 7 |
| (±)-geosmin | 4.259 ± 0.425 | -5.202 ± 0.064 | 0.816 ± 0.074 | 6 |
| heptanoic acid | 0.262 ± 0.173 | -2.000 ± 0.000 | 0.131 ± 0.086 | 5 |
| hexanal | 1.603 ± 0.164 | -2.000 ± 0.000 | 0.801 ± 0.082 | 5 |
| indole | 3.695 ± 0.547 | -4.328 ± 0.160 | 0.850 ± 0.117 | 6 |
| isobutyl acetate | 1.656 ± 0.364 | -2.000 ± 0.000 | 0.694 ± 0.059 | 6 |
| isopropyl tiglate | 2.953 ± 0.305 | -3.934 ± 0.219 | 0.758 ± 0.092 | 5 |
| L-(+)-lactic acid | 0.061 ± 0.008 | -2.000 ± 0.000 | 0.031 ± 0.004 | 5 |
| (*R*)-(+)-limonene | 4.302 ± 0.519 | -4.853 ± 0.099 | 0.882 ± 0.101 | 6 |
| linalool | 2.723 ± 0.268 | -3.996 ± 0.060 | 0.685 ± 0.074 | 6 |
| methyl benzoate | 2.810 ± 0.220 | -4.429 ± 0.178 | 0.631 ± 0.028 | 6 |
| methyl hexanoate | 3.961 ± 0.482 | -3.933 ± 0.043 | 1.005 ± 0.119 | 6 |
| methyl laurate | 0.054 ± 0.014 | -2.000 ± 0.000 | 0.027 ± 0.007 | 6 |
| o-cresol | $1.696 \pm 0.162$ | $-2.000 \pm 0.000$ | $0.848 \pm 0.081$ | 6 |
| octanoic acid | $0.071 \pm 0.012$ | $-2.000 \pm 0.000$ | $0.035 \pm 0.006$ | 5 |
| prenyl acetate | $1.614 \pm 0.193$ | $-2.000 \pm 0.000$ | $0.807 \pm 0.096$ | 6 |
| propyl acetate | $0.822 \pm 0.218$ | $-2.000 \pm 0.000$ | $0.411 \pm 0.109$ | 5 |
| sulcatone | $1.732 \pm 0.142$ | $-2.000 \pm 0.000$ | $0.866 \pm 0.071$ | 6 |
| thiazole | $0.207 \pm 0.064$ | $-2.000 \pm 0.000$ | $0.104 \pm 0.032$ | 5 |
| *trans*-3-hexen-1-ol | $0.260 \pm 0.026$ | $-2.000 \pm 0.000$ | $0.130 \pm 0.013$ | 5 |

**Extended Data Table 10.** Molecular descriptors of the ligands used in multiple regression analysis (Extended Data Fig. **13**). 1: Area, 2: Molecular weight (g/mol), 3: estimated octanol/water partition coefficient (XlogP3-AA), 4: Hydrogen bond count, donor, 5: Hydrogen bond count, acceptor, 6: Hydrogen bond count, total, 7: Rotatable bond count, 8: Polar surface area (Å^2^), 9: Water solubility (g/L), 10: Vapor pressure (mm Hg), 11: Polarizability (Å^3^). Sources: PubChem, Sigma-Aldrich, ChemSpider, EPA, and The Good Scents Company.

| Ligand | 1 | 2 | 3 | 4 | 5 | 6 | 7 | 8 | 9 | 10 | 11 |
| --- | --- | --- | --- | --- | --- | --- | --- | --- | --- | --- | --- |
| 1-hexanol | 151.07 | 102.17 | 2 | 1 | 1 | 2 | 4 | 20.2 | 5.9 | 0.93 | 12.4 |
| 1-octanol | 181.17 | 130.23 | 3 | 1 | 1 | 2 | 6 | 20.2 | 0.5 | 0.08 | 16.1 |
| (*R*)-1-octen-3-ol | 177.67 | 128.21 | 2.6 | 1 | 1 | 2 | 5 | 20.2 | 1.93 | 0.3 | 16.0 |
| 1-pentanol | 132.42 | 88.15 | 1.6 | 1 | 1 | 2 | 3 | 20.2 | 22 | 2.2 | 10.6 |
| 2-acetylthiophene | 134.58 | 126.18 | 1.2 | 0 | 2 | 2 | 1 | 45.3 | 14 | 3.72 | 13.7 |
| 2-ethylphenol | 147.69 | 122.16 | 2.5 | 1 | 1 | 2 | 1 | 20.2 | 3 | 0.15 | 14.9 |
| 2-heptanone | 161.14 | 114.19 | 2 | 0 | 1 | 1 | 4 | 17.1 | 4.3 | 3.85 | 13.7 |
| 2-undecanone | 219.98 | 170.29 | 4.1 | 0 | 1 | 1 | 8 | 17.1 | 0.02 | 0.04 | 21.0 |
| 2,3-butanediol | 117.17 | 90.12 | -0.9 | 2 | 2 | 4 | 1 | 40.5 | 1000 | 0.24 | 9.34 |
| 2,4,5-trimethylthiazole | 147.8 | 127.21 | 2.2 | 0 | 2 | 2 | 0 | 41.1 | 0.53 | 3.76 | 14.7 |
| 3-octanol | 117.17 | 130.23 | 2.8 | 1 | 1 | 2 | 5 | 20.2 | 1.3 | 0.51 | 16.1 |
| 4-ethylphenol | 149.23 | 122.16 | 2.6 | 1 | 1 | 2 | 1 | 20.2 | 7.4 | 0.04 | 14.9 |
| 4-methoxyphenylacetone | 189.56 | 164.2 | 1.6 | 0 | 2 | 2 | 3 | 26.3 | 3.08 | 0.008 | 18.7 |
| acetaldehyde diethyl acetal | 157.28 | 118.18 | 0.8 | 0 | 2 | 2 | 4 | 18.5 | 44 | 27.6 | 13.2 |
| acetophenone | 143.45 | 120.15 | 1.6 | 0 | 1 | 1 | 1 | 17.1 | 6.13 | 0.4 | 14.4 |
| alpha-pinene | 169.51 | 136.23 | 2.8 | 0 | 0 | 0 | 0 | 0 | 0.0025 | 4.75 | 17.4 |
| *L*-(+)-arabinose | 141.3 | 150.13 | -2.5 | 4 | 5 | 9 | 0 | 90.2 | 500 | 6.2 x 10<sup>-9</sup> | 12.5 |
| benzaldehyde | 126.34 | 106.12 | 1.5 | 0 | 1 | 1 | 1 | 17.1 | 6.95 | 1.27 | 13.1 |
| butyl acetate | 153.06 | 116.16 | 1.8 | 0 | 2 | 2 | 4 | 26.3 | 5.3 | 11 | 12.5 |
| butyric acid | 111.45 | 88.11 | 0.8 | 1 | 2 | 3 | 2 | 37.3 | 60 | 1.65 | 8.78 |
| caffeine | 187.53 | 194.19 | -0.1 | 0 | 3 | 3 | 0 | 58.4 | 21.6 | 9 x 10<sup>-7</sup> | 20.0 |
| decanal | 204.92 | 156.26 | 3.8 | 0 | 1 | 1 | 8 | 17.1 | 0.029 | 0.1 | 19.3 |
| DEET | 220.87 | 191.27 | 2 | 0 | 1 | 1 | 3 | 20.3 | 0.91 | 0.002 | 23.3 |
| ethyl acetate | 117.6 | 88.11 | 0.7 | 0 | 2 | 2 | 2 | 26.3 | 80 | 93.2 | 8.86 |
| ethyl butyrate | 155.12 | 116.16 | 1.3 | 0 | 2 | 2 | 4 | 26.3 | 4.9 | 14 | 12.5 |
| ethyl hexanoate | 188.21 | 144.21 | 2.4 | 0 | 2 | 2 | 6 | 26.3 | 0.63 | 1.78 | 16.2 |
| eugenol | 189.2 | 164.2 | 2 | 1 | 2 | 3 | 3 | 29.5 | 2.46 | 0.02 | 19.3 |
| (±)-geosmin | 196.36 | 182.3 | 3.3 | 1 | 1 | 2 | 0 | 20.2 | 0.157 | 0.003 | 21.8 |
| D-(+)-glucose | 165.56 | 180.16 | -2.6 | 5 | 6 | 11 | 1 | 110 | 960 | 5.9 x 10<sup>-11</sup> | 14.9 |
| heptanoic acid | 164.52 | 130.18 | 2.5 | 1 | 2 | 3 | 5 | 37.3 | 2.82 | 0.01 | 14.3 |
| hexanal | 143.01 | 100.16 | 1.8 | 0 | 1 | 1 | 4 | 17.1 | 5.64 | 11.3 | 11.9 |
| indole | 134.94 | 117.15 | 2.1 | 1 | 0 | 1 | 0 | 15.8 | 3.56 | 0.01 | 15.3 |
| isobutyl acetate | 189.2 | 116.16 | 1.8 | 0 | 2 | 2 | 3 | 26.3 | 6.3 | 17.8 | 12.5 |
| isopropyl tiglate | 180.68 | 142.2 | 2.1 | 0 | 2 | 2 | 3 | 26.3 | 0.5 | 3.51 | 16.2 |
| *L*-(+)-lactic acid | 101.64 | 90.08 | -0.7 | 2 | 3 | 5 | 1 | 57.5 | 1000 | 0.08 | 7.53 |
| (*R*)-(+)-limonene | 179.13 | 136.25 | 3.4 | 0 | 0 | 0 | 1 | 0 | 0.007 | 1.64 | 18.0 |
| linalool | 200.28 | 154.25 | 2.7 | 1 | 1 | 2 | 4 | 20.2 | 1.59 | 0.16 | 19.6 |
| methyl benzoate | 154.73 | 136.15 | 2.1 | 0 | 2 | 2 | 2 | 26.3 | 2.1 | 0.38 | 15.1 |
| methyl hexanoate | 170.24 | 130.18 | 2.5 | 0 | 2 | 2 | 5 | 26.3 | 1.33 | 3.72 | 14.4 |
| methyl laurate | 278.56 | 214.34 | 5.8 | 0 | 2 | 2 | 11 | 26.3 | 0.013 | 4.11x10<sup>-3</sup> | 25.4 |
| *n*-caproic acid | 147.04 | 116.16 | 1.9 | 1 | 2 | 3 | 4 | 37.3 | 10.3 | 0.04 | 12.5 |
| *o*-cresol | 130.17 | 108.14 | 2 | 1 | 1 | 2 | 0 | 20.2 | 25.9 | 0.3 | 13.1 |
| octanoic acid | 180.9 | 144.21 | 3 | 1 | 2 | 3 | 6 | 37.3 | 0.789 | 3.71x10<sup>-3</sup> | 16.1 |
| prenyl acetate | 164.81 | 128.17 | 1.8 | 0 | 2 | 2 | 3 | 26.3 | 3.2 | 5.4 | 14.3 |
| propyl acetate | 138.16 | 102.13 | 1.2 | 0 | 2 | 2 | 3 | 26.3 | 18.9 | 35.9 | 10.7 |
| sulcatone | 171.92 | 126.2 | 1.9 | 0 | 1 | 1 | 3 | 17.1 | 3.2 | 0.008 | 15.5 |
| thiazole | 90.71 | 85.13 | 0.4 | 0 | 2 | 2 | 0 | 41.1 | 53 | 17.15 | 9.01 |
| *trans*-3-hexen-1-ol | 144.57 | 100.16 | 1.3 | 1 | 1 | 2 | 3 | 20.2 | 16 | 1.36 | 12.5 |

## Notes

### Competing Interest Statement

The authors have declared no competing interest.

